# Dps binds and protects DNA in starved *Escherichia coli* with minimal effect on chromosome accessibility, dynamics and organisation

**DOI:** 10.1101/2025.08.31.673347

**Authors:** Lauren A. McCarthy, Lindsey E. Way, Xiaofeng Dai, Zhongqing Ren, David E. H. Fuller, Ishika Dhiman, Linaria Larkin, Jurriaan J. D. Sieben, Ilja Westerlaken, Elio A. Abbondanzieri, Gail G. Hardy, Anne S. Meyer, Xindan Wang, Julie S. Biteen

## Abstract

Dps is the most abundant nucleoid-associated protein in starved *Escherichia coli* with ∼180,000 copies per cell. Dps binds DNA and oxidises iron, facilitating survival in harsh environments. Dps-DNA complexes can form crystalline structures, leading to the proposed model that Dps reorganises the starved *E. coli* nucleoid into a compact liquid crystal, slowing chromosome dynamics and limiting access of other proteins to DNA. In this work, we directly tested this model using live-cell super-resolution microscopy and Hi-C analysis. We found that after 96 h of starvation, Dps compacts the nucleoid and increases short-range DNA-DNA interactions, but does not affect chromosome accessibility to large protein nanocages or small restriction enzymes. We also report that chromosome dynamics and organisation are primarily impacted by the bacterial growth phase; the effect of Dps is relatively minor. Our work clarifies the role of Dps in modulating nucleoid properties, and we propose an updated model for Dps-DNA interactions in which Dps binds, protects and compacts DNA largely without influencing chromosome access, dynamics and organisation. Additionally, this work provides a general framework for assessing the impact of nucleoid-associated proteins on key aspects of chromosome function in live cells.

**TOC:** 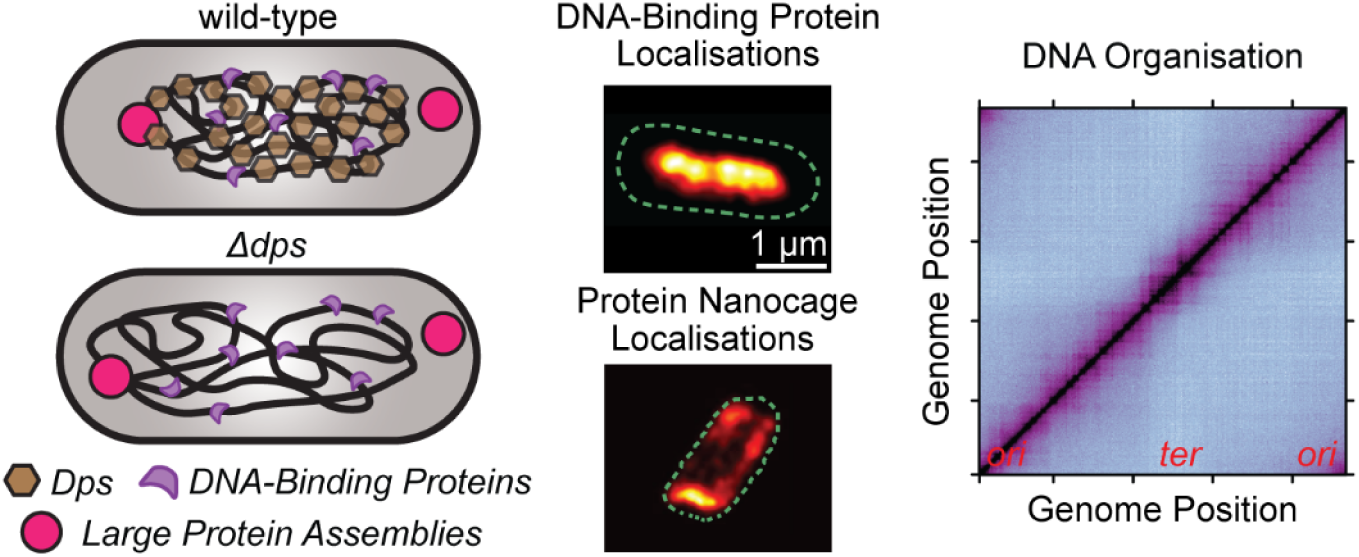

## INTRODUCTION

Bacterial cells lack a membrane-bound nucleus and eukaryotic histones and instead use alternative methods to compartmentalise, protect and organise genomic DNA within the bacterial nucleoid (1, 2). The DNA-binding protein from starved cells (Dps) is a conserved protein across the bacterial kingdom that protects DNA from many stressors, including oxidative damage, acidic and basic shock, UV radiation and antibiotic exposure (3–5). In *Escherichia coli*, Dps responds to various stressors through dual functionality (4, 6). Dps binds DNA, providing physical protection from harmful species (4, 7, 8), and Dps processes hydrogen peroxide to store iron safely, preventing DNA-damaging radicals from forming (6, 9, 10). Dps is highly expressed in starved cells, facilitating the survival of *E. coli* in harsh, nutrient-poor environments (11).

Dps belongs to the family of nucleoid-associated proteins (NAPs) in *E. coli* and is the most abundant NAP in stationary-phase (i.e., starved) cells with ∼180,000 copies after 48 h of incubation (12). NAPs have garnered attention for their essential role in regulating the architecture and organisation of the bacterial nucleoid (1, 13, 14). Dps typically oligomerises into dodecamers (15, 16), giving each Dps oligomer multiple binding sites to DNA and other Dps oligomers (17). These highly cooperative interactions (18, 19) condense the bacterial nucleoid in stationary phase (7, 20), which is expected to affect nucleoid organisation. However, the morphology of Dps-condensed nucleoids and the resulting impact on the spatial arrangement, accessibility and conformation of the chromosome remain poorly understood.

Electron microscopy (EM) images of Dps-condensed nucleoids in *E. coli* reveal striking arrays of Dps and DNA, suggesting that the Dps can reorganise the chromosome into a hexagonally packed biocrystal (21). These results lead to the commonly accepted model that Dps dramatically restructures the nucleoid in stationary phase (**Figure 1**). EM has also indicated other possible Dps-DNA structures, including toroids in starved *E. coli* cells (22) that are more likely to occur early in stationary phase. However, EM is often performed after cell fixation, and many of these assays were carried out in cells artificially overexpressing Dps. Therefore, it is unclear how the nucleoid is restructured by Dps in the wild-type (*wt*), live-cell context. On the other hand, Janissen *et al*. demonstrated that transcription is largely unaffected by Dps and that Dps-DNA complexes are dynamic, allowing RNA polymerase to quickly access and bind Dps-coated DNA (7). The same study also found that *in vitro*, Dps protects DNA from digestion by restriction enzymes. The selective protection offered by Dps suggests that Dps-DNA complexes form dynamic liquid crystalline structures that are selectively soluble for some proteins but not others (7). Based on these results, the current model for DNA protection and organisation by Dps (**Figure 1**) indicates that: *(a)* starvation induces high Dps expression, which condenses and restructures the chromosome into a dynamic liquid crystal (3, 7, 23); *(b)* size-based steric hindrance excludes large protein assemblies from the Dps-condensed nucleoid (3, 24); *(c)* DNA binding proteins can either be included or excluded from the nucleoid based on selective solubility, possibly related to protein function (7); and *(d)* in cells lacking Dps, the nucleoid is less condensed, lacks liquid crystalline structure and is accessible to small and large proteins.

**Figure 1.**
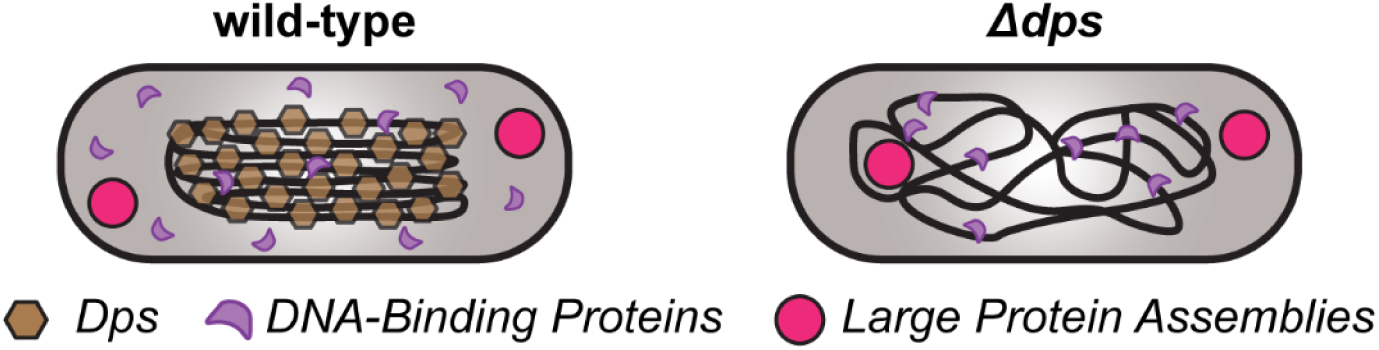
Current model for starvation-induced nucleoid protection and organisation mediated by Dps.

In this study, we analysed Dps-mediated nucleoid condensation and organisation by conducting multiscale assays in living cells, both with and without Dps (*Δdps*). First, we characterised Dps-dependent nucleoid compaction with super-resolution optical microscopy. Next, we examined whether Dps modulates the nucleoid accessibility to a large nanoparticle (protein nanocage assembly), a small restriction enzyme (EcoRI) and RNA polymerase. We then characterised the dynamics and spatial arrangement of the chromosome with single-particle tracking. Finally, we quantified the role of Dps in maintaining short-range and long-range DNA-DNA interactions within the genome by combining chromosome-conformation capture techniques with next-generation sequencing (Hi-C). Our analyses reveal that, despite high Dps expression in stationary phases, Dps has a relatively minor effect on chromosome accessibility, dynamics and organisation. Instead, Dps is able to bind and protect DNA, measurably compact the nucleoid after 96 h of starvation and increase short-range DNA-DNA interactions, all with minimal interference with other nucleoid properties. Additionally, we find that in both *wt* and *Δdps* cells, chromosome dynamics and organisation depend strongly on the growth phase.

## MATERIAL AND METHODS

### *E. coli* strains and growth conditions

All strains used in this study were derived from prototrophic *E. coli* K-12 strain W3110 (CGSC no. 4474) (25). Cells were streaked from glycerol stocks onto LB-agar plates containing antibiotics when appropriate at the following concentrations: 20 µg/mL kanamycin (CAS no. 25389-94-0), 20 µg/mL chloramphenicol (CAS no. 56-75-7), or 100 µg/mL spectinomycin (CAS no. 22189-32-8). Single colonies were then isolated from plates and grown in liquid culture. Unless noted otherwise, liquid cultures were grown in a High-Def Azure (HDA) medium (Teknova cat. No. 3H5000) supplemented with 0.2% glucose (m/v), which, for simplicity, is referred to as “HDA medium” throughout the manuscript. Cultures were incubated at either 30 °C or 37 °C (as indicated) with shaking at 250 rev/min. The strains, plasmids, oligonucleotides and Next-Generation-Sequencing samples are listed in Tables S1 – S4.

### Fluorescence Microscopy

#### Photoactivated Localisation Microscopy (PALM) and Single-Molecule Tracking

PALM imaging and tracking were performed by activating a few single molecules of PAmCherry at a time, imaging those molecules until they bleached, activating a new set of molecules and then repeating this imaging process over several minutes. PALM experiments were completed on an Olympus IX-71 microscope with either a 100× 1.40 NA or a 100× 1.45 NA phase-contrast, oil-immersion objective heated to 30 °C with an objective heater (Bioptics). Immersion oil optimised for 30 °C (Zeiss) was used to reduce optical aberrations. Photoactivation was performed with a 406-nm laser (Coherent Cube 405-100) with a 0.5 – 2 W/cm^2^ power density and exposure times optimised for each construct (100 – 400 ms). PAmCherry fusions were imaged with a 561-nm laser (Coherent Sapphire 561-50) with power densities of 0.46 and 0.55 kW/cm^2^ for HUα-PAmCherry and PAmCherry-EcoRI[E111Q], respectively. Fluorescence emission from PAmCherry was filtered with a 561-nm long-pass filter and imaged with a 512 × 512-pixel Photometrics Evolve electron-multiplying charge-coupled detector (EMCCD) camera. Each cell was imaged for no longer than 6.5 min to prevent phototoxicity. Each prepared sample of cells on an agarose pad was imaged for no more than 1 h.

Cells expressing HUα-PAmCherry from the native promoter were grown overnight at 30 °C with shaking at 250 rpm in HDA medium and then diluted 1:100 into 50 mL of fresh HDA medium followed by continued incubation at 30 °C for 24 h or 96 h. Single colonies of cells encoding *pamCherry-ecoRI[E111Q]* on an arabinose-inducible promoter on plasmid pLAM016 were obtained from freshly streaked LB-agar plates supplemented with 100 µg/mL spectinomycin and 0.2% glucose, which was added to suppress expression. Overnight cultures were grown at 30 °C in HDA medium with spectinomycin. Saturated overnight cultures were back-diluted to OD_600_ = 0.03 – 0.05 in 25 – 50 mL of fresh HDA medium with spectinomycin and 0.025% arabinose and allowed to incubate at 30 °C for ∼96 h.

After the appropriate incubation time, spent medium was prepared by centrifuging 10 mL of saturated culture from each culture flask for 6.5 – 7.5 min at 4950 – 6600 × *g* and syringe filtering the supernatant at least twice using a fresh 0.22-µm syringe filter each time. Spent medium was then used to make 2% (m/v) agarose pads, and 1.5 µL of as-grown cells were deposited onto the agarose pad along with 1.5 µL of Fluoresbrite carboxylate YG beads (0.35 µm, Polysciences) that had been previously suspended in spent medium (1 – 3 µL as-shipped beads diluted into 1 mL of spent medium). The cells and beads were sandwiched onto the agarose pad by a No. 1 coverslip.

#### Protein Nanocage Imaging and Tracking

eGFP-labelled protein nanocages (26) were expressed from an arabinose-inducible promoter on plasmid pLAM003. For the 24-h time point, 30 µL of saturated overnight culture grown at 30 °C was diluted into 3 mL of fresh HDA media containing 100 µg/mL spectinomycin and 8 – 12 µg/mL arabinose, and the culture was incubated for 24 h at 30 °C. For the 96-h time point, 200 µL of saturated overnight culture grown at 37 °C was diluted into 20 mL of fresh HDA medium containing 100 µg/mL spectinomycin and 18 – 20 µg/mL arabinose, and the culture was incubated for 96 h at 30 °C.

After the elapsed incubation time, spent medium was prepared by centrifuging culture and filtering the supernatant twice as described above. Then cells were concentrated by 10×, and 2 µL of concentrated cells was pipetted directly onto a 2% (w/v) agarose pad made from spent medium and sandwiched by a No. 1 coverslip.

Imaging was performed on an Olympus IX-71 inverted microscope with a 100× 1.40 NA oil-immersion objective heated to 30°C by an objective heater (Bioptics) and using appropriate index-matched oil. The cell sample was mounted on the microscope objective and allowed to rest for 20 min to thermally equilibrate before imaging. Each sample was imaged for 60 – 75 min. Fluorescence imaging of the nanocage-eGFP fusions was performed with a 488-nm laser (Coherent Sapphire 488-550) with a power density of 100 W/cm^2^. Movies capturing nanocage diffusion were acquired continuously with a 512 × 512-pixel Photometrics Evolve EMCCD camera (40-ms frame time).

#### Chromosomal Locus Imaging and Tracking

Fluorescence microscopy of the loci using continuous imaging with a short exposure time (40 ms) was performed with the same microscope and camera described above for PALM imaging. Excitation was from a 488-nm laser (Coherent-Sapphire 488-50) with a power density of 200 W/cm^2^, and appropriate optical filters were used to image the emission from YGFP. Cells were grown overnight in Luria Broth medium (Fisher) supplemented with 20 µg/mL of kanamycin at 37 °C for 12 – 16 h and then were sub-cultured (1:100 dilution) in pre-warmed HDA medium and incubated at 30 °C. For imaging cells after 24 h of incubation, two subcultures were prepared: a 3-mL culture for microscopy and a 5-mL culture to prepare spent medium. After 24 h of culturing, the paired 24-h culture was centrifuged and filtered twice with a 0.22-µm filter to prepare spent medium, which was then used to make 2% agarose pads. Cells were then immobilised on the agarose pads and imaged with an objective that was heated to 30 °C using an objective heater (Bioptics) and oil that was index-matched to 30 °C. For imaging cells after 96 h of incubation, precultures were diluted (1:100) into 50 mL of HDA medium. After 96 h, 5 mL of culture was used to prepare spent medium for making 2% agarose pads as described above. All other imaging procedures described for 24 h were repeated at 96 h.

Fluorescence microscopy of the loci using a 2-s time interval was performed with a Nikon Instruments Ti2 microscope equipped with a Plan Apo 100× 1.45 NA phase-contrast oil-immersion objective, a SOLA SE II 365 Light Engine as light source, a Nikon C-FLL Large Field of View YFP filter set and a Hamamatsu ORCA-Flash 4.0 LT+ sCMOS camera. Cells were grown in HDA medium at 37 °C for approximately 5 h and then were subcultured in 30 mL of pre-warmed HDA medium to an initial OD_600_ of 0.03. For each strain, three subcultures were prepared: one for microscopy samples, one to prepare spent medium at 24 h and one for spent medium at 96 h. Exponential-phase cells were collected at OD_600_ 0.3 – 0.4 and immobilised on 2% (w/v) agarose pads containing fresh HDA medium. Spent medium for the 24-h time point was prepared after 23.5 h of culture growth; the culture was centrifuged and filtered twice: once with a 0.45-µm filter and once with a 0.20-µm filter. This spent medium was used to prepare 2% agarose pads. After 24 h of culture growth, cells were collected and immobilised for imaging on these agarose pads. The process for the 24-h time point was repeated at 96 h. Images and videos were captured using Nikon NIS-Elements software.

### Fluorescence Microscopy Data Analysis

#### Single-Molecule and Single-Particle Localisation and Tracking

Cell segmentation from phase-contrast cell images was performed with the publicly available Cellpose (27) or Omnipose (28) packages in Python that were trained on bacterial data. Only data within a cell outline was analysed. Detection, localisation and tracking of single fluorescent molecules, particles and foci were performed with the SMALL-LABS algorithm (29).

#### Calculation of Localisation Heatmaps

For each segmented cell, the length (long axis) and width (short axis) were calculated from the maximum and minimum Feret diameters, respectively. Cells with lengths 1.5 µm – 4.0 µm were chosen for further analysis. This cell length filter captures >90% of cellular morphologies (**Figure S1**). The coordinates of the fluorescent targets within each cell were normalised by the corresponding length of the long and short axes of the cell. A 2D histogram was used to visualise the ensemble localisation density of the normalised coordinates of the target molecules; these histograms are referred to as “heatmaps” throughout the manuscript. The heatmaps are symmetrised along the long and short axes and normalised such that the sum of all pixel intensities is one. The aspect ratio used to display the heatmap for each condition indicates the average aspect ratio for each condition, which was determined by applying the cell-length filter to cell dimensions sampled from HUα-PAmCherry *wt* and *Δdps* cells measured at 24 h and 96 h of starvation (**Table 1**). All heatmaps at the same condition were rendered with this aspect ratio. To ensure that only localisations with the highest precision were used to determine the nucleoid outlines in Figure 3 originating from the HUα-PAmCherry heatmaps, only localisations with a 95% confidence interval <80 nm were used to calculate the HUα-PAmCherry heatmaps in Figure 2.

**Figure 2.**
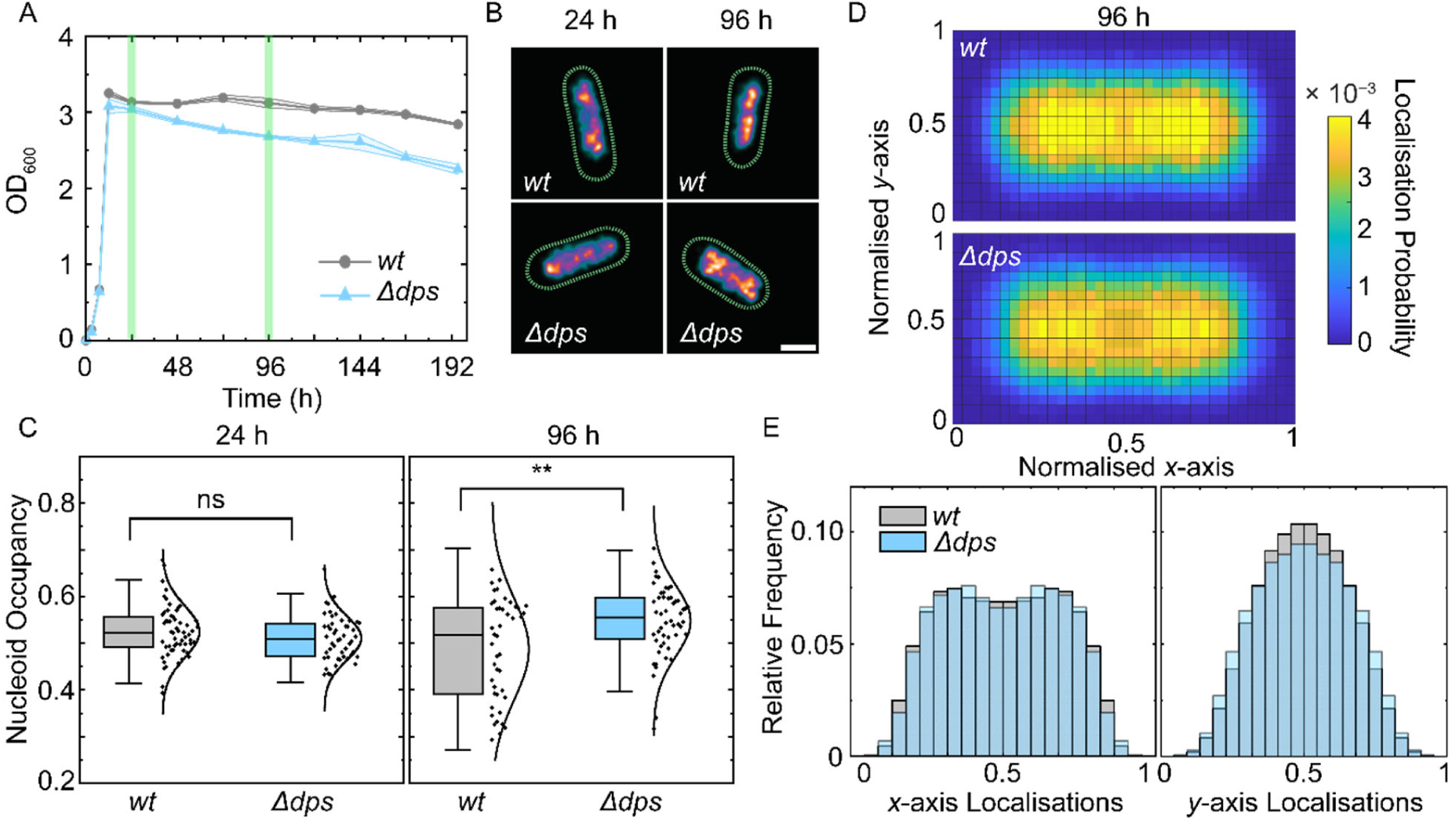
Live-cell super-resolution imaging of nucleoid occupancy and HUα-PAmCherry spatial distributions. **(A)** *wt* and Δ*dps* HUα-PAmCherry cell growth curves; shading is the standard deviation from three biological replicates. The green lines mark the time points of interest to this study: stationary phase (24 h) and deep stationary phase (96 h). **(B)** Representative super-resolution images of *wt* and Δ*dps* HUα-PAmCherry *E. coli* at each time point, constructed from 1500 – 2000 localisations/cell, each with 95% confidence interval ≤ 80 nm. Colour corresponds to the localisation density: high densities are orange and low densities are blue. **(C)** Nucleoid occupancy distributions for *wt* and Δ*dps* cells in stationary phase (left) and deep stationary phase (right) with *n* = 44 – 58 cells per condition. The whiskers show 2 standard deviations from the mean, and the centre line is the median. Distributions of nucleoid occupancies are marked as either not significantly different (ns, *p* > 0.05) or with stars (**, *p* < 0.01) from a Kolmogorov-Smirnov test. **(D)** Normalised heatmaps of HUα-PAmCherry localisations in deep stationary phase for *wt* and Δ*dps* cells. The box outline was rendered with the same aspect ratio as the average cell for each population. **(E)** Histograms of the *x*-axis and *y*-axis localisation probabilities from the heatmaps in **D**.

**Figure 3.**
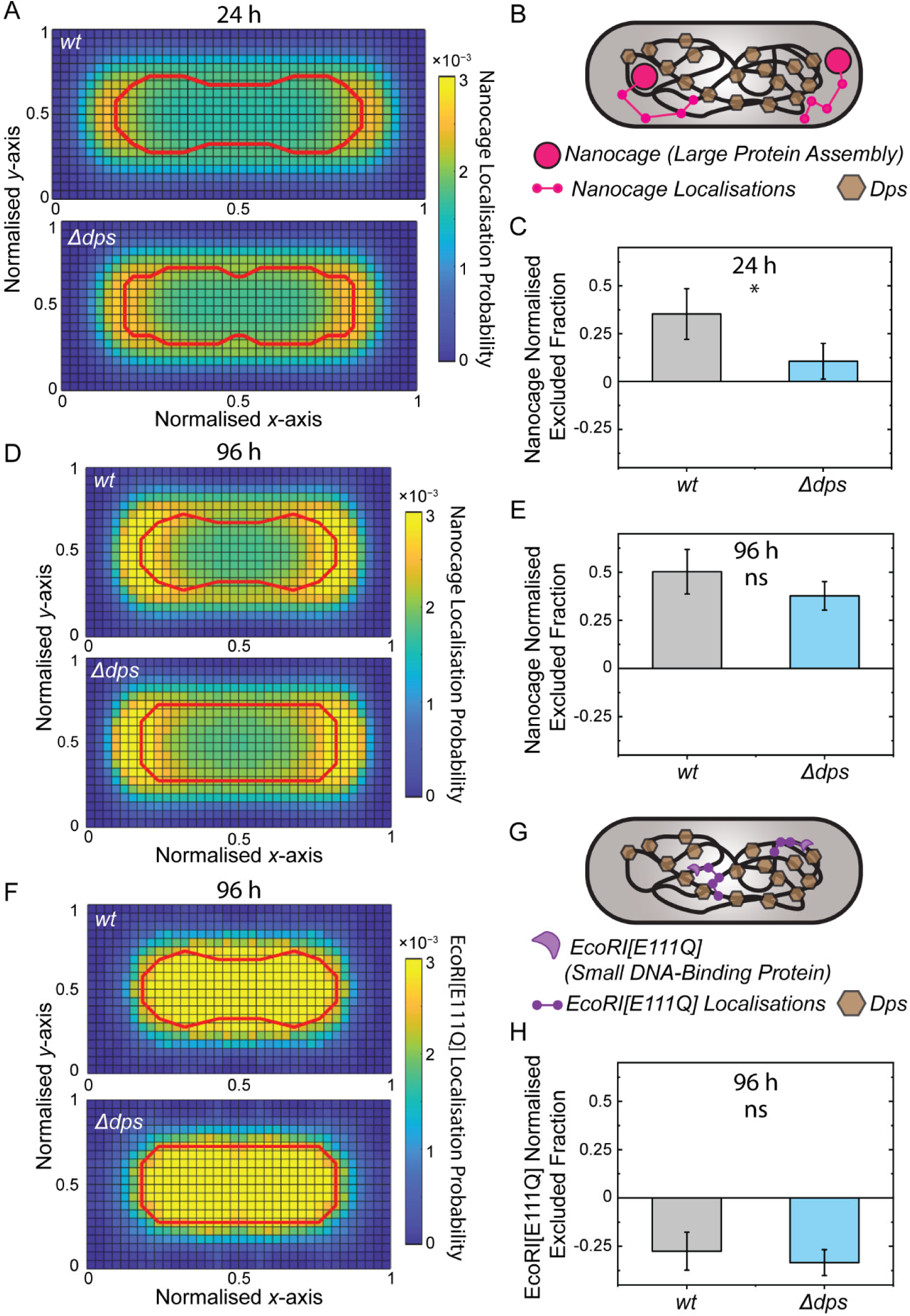
Exclusion from the nucleoid of probes of different sizes in cells with and without Dps. **(A)** Heatmap of nanocage localisations for *wt* and *Δdps* cells in stationary phase. The red outlines display the nucleoid boundaries calculated from the top 60% contour line of localisation probabilities in the HUα-PAmCherry heatmaps for *wt* and *Δdps* cells measured at 24 h of starvation (**Figure S4**). **(B)** Illustration of a nanocage (pink dots) diffusing in a *wt E. coli* cell. **(C)** Normalised excluded fraction from the nucleoid for nanocages in stationary phase. The height of the bar corresponds to the mean from five randomly sampled data sets (without replacement) of ∼800 cells collected from two biological replicates. Error bars: standard deviation of the five sub-sampled data sets. *: statistically different (*p* < 0.05) in a two-sample t-test. **(D, E)** Same as **(A, C)** but for cells in deep stationary phase. The red outlines in **(D)** display the nucleoid outlines calculated as described from the HUα-PAmCherry heatmaps for *wt* and *Δdps* cells measured at 96 h (Figure 1D). “ns” in **(E)** indicates that the means are not significantly different (*p* > 0.05) by a two-sample t-test. **(F)** Same as **(D)** but for drift-corrected PAmCherry-EcoRI[E111Q] localisations, with drift correction performed every 16 – 40 s. To ensure that only cells with adequate PAmCherry-EcoRI[E111Q] expression were considered, cells with < 100 localisations were excluded from the analysis. **(G)** Cartoon of PAmCherry-EcoRI[E111Q] (purple dots) diffusing within a *wt E. coli* cell. **(H)** Normalised excluded fraction from the nucleoid for PAmCherry-EcoRI[E111Q] molecules in deep stationary phase. The height of the bar corresponds to the mean from three randomly sampled data sets (without replacement) of ∼10 cells each collected from four biological replicates. Error bars: standard deviation from the three sub-sampled data sets.

**Table 1.** Calculated aspect ratios for *wt* and *Δdps E. coli* cells at each growth condition.

| | *wt* | $\Delta dps$ |
| --- | --- | --- |
| Average aspect ratio after 24 h of growth at 30 °C | 2.2:1 | 2.1:1 |
| Average aspect ratio after 96 h of growth at 30 °C | 1.8:1 | 1.8:1 |

#### Reconstruction of Nucleoid PALM Images and Calculation of Nucleoid Occupancy

During PALM imaging of HUα-PAmCherry, sample drift was monitored with a fluorescent bead near the cells. We tracked this drift within the 3-min videos by determining the average bead position every 8 s based on fitting the fluorescence intensity from 200 summed imaging frames to a 2D Gaussian. The molecule positions were then updated based on the corresponding drift for each frame number of each single-molecule localisation.

To reconstruct PALM images of the nucleoid, all drift-corrected HUα-PAmCherry localisations with 95% confidence interval <80 nm were selected. A Gaussian kernel was applied to each localisation with the degree of blurring (*σ*) set to the confidence interval of that localisation.

To determine the nucleoid occupancy from the PALM images, the distribution of the pixel intensities was fit with a two-component Gaussian Mixture Model. Pixel intensities below the fifth percentile and above the 99.95 percentile were omitted to remove outliers. The threshold pixel value was set to 50% of the mean intensity of the second Gaussian distribution. Pixels with intensity greater than the threshold were considered to belong to the nucleoid region; the nucleoid occupancy was determined from the number of pixels in this nucleoid region divided by the number of pixels within the cell outline determined by Omnipose (28).

#### Calculation of the Fraction of Molecules Excluded from the Nucleoid

The nucleoid boundary was determined by the 60% contour line from the Huα-PAmCherry heatmap at each condition, and the apparent excluded fraction, *exc_app_*, is the integral of the normalised protein nanocage or EcoRI-PAmCherry occupancy heatmap outside of this nucleoid boundary. However, this 2D projection miscounts the excluded fraction because particles above and below the nucleoid are counted as inside the nucleoid boundary. To adjust for the 3D cell geometry, we transformed the 2D cell and nucleoid contours from the heatmap into 3D based on assuming radial symmetry and rotating about the cell long axis. We then projected a 3D cell with all particles in the cytoplasm (no particles in the nucleoid) back to 2D to determine the maximum apparent excluded fraction, *exc_max_* ; and we projected a 3D cell with an equal concentration of particles in the cytoplasm and nucleoid back to 2D to determine the corresponding excluded fraction, *exc*_0_.

The corrected, normalised excluded fraction, *exc_norm_*, was therefore calculated by comparing the apparent excluded fraction, *exc_app_*, to *exc_max_* and *exc*_0_:

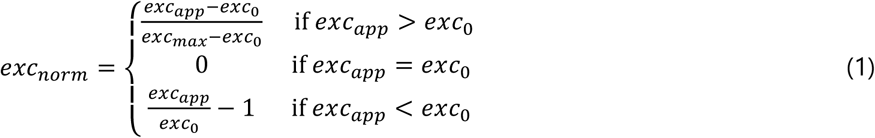

This normalisation process yields −1 ≤ *exc_norm_* ≤ 1, such that *exc_norm_* = −1 indicates that all particles are within the nucleoid, and *exc_norm_* = 1 indicates that all particles are nucleoid-excluded.

#### Diffusion Analysis

To quantify the trajectories of chromosomal loci from time-lapse videos acquired with a 2-s exposure time, drift correction was performed based on the simultaneously recorded phase contrast channel. A region at the centre of the image with sparse cell density was selected for a cross-correlation calculation between adjacent frames. Then, 2D Gaussian fitting was performed on the cross-correlation matrix to calculate the drift between each frame with subpixel precision. A cell length filter of 1.5 – 4.0 µm was applied. Next, to remove spurious localisations due to noise or debris, the DBSCAN algorithm (30) was applied to the chromosomal locus localisations to identify clustered regions. Only clusters detected in more than 8 frames were used for further analysis and identified as chromosomal loci. To ensure that cells with similar DNA content were compared, cells with one focus were analysed separately from those with two foci.

After correcting for drift and filtering the cells, the diffusion coefficient was calculated by mean-squared displacement (MSD) analysis (31). For each trajectory, only the first 30% of displacements were used for curve fitting, which is the range corresponding to a linear relationship between time lag and displacement despite the overall sub-diffusive motion of the loci. Linear fitting was used to calculate the diffusion coefficient, and results with *R*^2^ < 0.7 were removed from further consideration. The distributions of diffusion coefficients were used to quantify the mobility of loci from each specific condition.

To calculate the confinement area for each locus, the *boundary* function in Matlab was applied to each set of cluster localisations determined by DBSCAN. The area of the polygon corresponding to the detected boundary was assigned as the confinement area.

### Whole Genome Sequencing

Whole genome sequencing was performed as previously described (32). Briefly, cells were grown to the desired growth phase in HDA medium at 37 °C. Genomic DNA was extracted using the Qiagen DNeasy blood and tissue kit (Qiagen, 69504). DNA was sonicated using a Qsonica Q800R2 sonicator for 12 min at 20% amplitude to achieve an average fragment size of 170 bp. The DNA library was prepared using NEBNext Ultra II kit (E7645) and sequenced using Illumina NextSeq500 or Nextseq2000 at the Indiana University Center for Genomics and Bioinformatics. Sequencing reads were mapped to the *E. coli* W3110 genome (NCBI reference sequence GCA_000005845.2 with a deletion from 1398953 to 1438196) using CLC Genomics Workbench (Qiagen). The mapped reads were normalised to the total number of reads and used as input for the ChIP samples.

### ChIP-seq

Chromatin immunoprecipitation (ChIP) was performed as previously described (33). Cells were grown to the desired growth phase in HDA medium at 37 °C. Cells were crosslinked in 3% formaldehyde for 30 min at room temperature. The cells were quenched using 125 mM glycine, washed with PBS and lysed using lysozyme. A Qsonica Q800R2 sonicator was used to shear crosslinked chromatin to an average size of 170 bp. The lysate was precleared using Protein A magnetic beads (GE Healthcare/Cytiva 28951378) and then incubated with anti-Dps antibodies (34) overnight at 4 °C. The following day, the lysate was incubated with Protein A magnetic beads for 1 h at 4 °C. After washes and elution, the immunoprecipitates were incubated at 65 °C overnight to reverse the crosslinking. The DNA was further treated with RNaseA, Proteinase K, extracted with a phenol/chloroform/isoamylalcohol (PCI) mixture (25:24:1), resuspended in 100 µL of Qiagen EB buffer, and used for library preparation with the NEBNext Ultra II kit (E7645). Library sequencing was performed using Illumina NextSeq500 or NextSeq2000 at the Indiana University Center for Genomics and Bioinformatics. The sequencing reads were mapped to the genome of *E. coli* W3110 (NCBI reference sequence GCA_000005845.2 with a deletion from 1398953 to 1438196) using CLC Genomics Workbench (Qiagen). Sequencing reads were normalised by the total number of reads, and plotted and analysed using R. The *x*-axis was rearranged to start at the replication origin, with the terminus region near the middle of the plot.

### Hi-C DNA-DNA Interaction Measurements

Cells grown to the desired condition were crosslinked with 10% formaldehyde at room temperature for 30 min then quenched with 125 mM glycine. Cells were lysed using Ready-Lyse Lysozyme (Epicentre, R1802M) and treated by 0.5% SDS. Solubilised chromatin was digested with Sau3AI for 2 h at 37 °C. The digested ends were filled in with Klenow and Biotin-14-dATP, dGTP, dCTP, dTTP. The products were ligated with T4 DNA ligase at 16 °C for about 20 h. Crosslinks were reversed at 65 °C for about 20 h in the presence of EDTA, proteinase K and 0.5% SDS. The DNA was then extracted twice with PCI, precipitated with ethanol, and resuspended in 20 µL of 0.1× TE buffer. Biotin from non-ligated ends was removed using T4 polymerase (4 h at 20 °C) followed by extraction with PCI. The DNA was then sheared by sonication for 12 min with 20% amplitude using a Qsonica Q800R2 sonicator. The sheared DNA was used for library preparation with the NEBNext UltraII kit (E7645). Biotinylated DNA fragments were purified using 10 µL streptavidin beads. DNA-bound beads were used for polymerase chain reaction (PCR) in a 50 µL reaction for 14 cycles. PCR products were purified using Ampure beads (Beckman, A63881) and sequenced at the Indiana University Center for Genomics and Bioinformatics using Illumina NextSeq500 or NextSeq2000. Paired-end sequencing reads were mapped to the genome of *E. coli* W3110 (NCBI reference sequence GCA_000005845.2 with a deletion from 1398953 to 1438196) using the same pipeline described previously (35). The *E. coli* W3110 genome was first divided into 461 10-kb bins. Subsequent analysis and visualisation were performed using R scripts. For the log_2_ ratio plots, the Hi-C matrix of strain 1 was divided by the matrix of strain 2. Then, log_2_(strain 1/strain 2) was calculated and plotted in a heatmap using R. The *x* and *y* axes were rearranged to start at the replication origin, with the terminus region near the centre of the map.

### Strain and Plasmid Construction

Plasmids were constructed using isothermal assembly of DNA fragments and strains were constructed using lambda recombineering. Complete details are in the Supplementary Material.

## RESULTS

### Dps increases the optical density of stationary-phase cultures

To assess the protective effect of Dps in stationary phase, we characterised the growth curves of *E. coli wt* and Δ*dps* cultures after inoculation into fresh HDA medium at 30 °C (**Figure 2A**). We found that the absence of Dps decreases the OD_600_ during stationary phase: *wt* cell cultures maintain a steady OD_600_ until 96 h, while in Δ*dps* cells, the OD_600_ steadily decreases after 24 h, suggesting an earlier onset to the death phase for *Δdps* cells. Given that Dps expression dramatically increases upon entry to stationary phase (∼13 h) (12), timepoints beyond 13 h can be used to study the impacts of Dps on chromosome compaction, accessibility, dynamics and organisation. However, to isolate biologically relevant impacts, we studied *wt* and *Δdps* cells after 24 and 96 h (green lines, **Figure 2A**). We refer to these time points throughout the manuscript as the “stationary phase” and “deep stationary phase,” respectively. At 24 h, *Δdps* and *wt* cultures have similar OD_600_, consistent with previous work assessing cell growth via colony-forming units (CFUs) after starvation (4). After 96 h, *wt* cell cultures exhibit an OD_600_ that is 0.35 greater than that of *Δdps* cells, consistent with CFU quantification after 48 h of starvation (4). Therefore, Dps has a protective effect in deep stationary phase, and any observed changes to nucleoid accessibility, dynamics and interactions induced by Dps at this time point are likely beneficial.

### Dps compacts the nucleoid in deep stationary phase

Consistent with previous work, we determined that Dps does not influence cell size in stationary and deep stationary phases (**Figure S1**) (7). We characterised nucleoid size in live *wt* and *Δdps* cells using a nucleic acid stain. Surprisingly, we found that Dps does not have a measurable impact on nucleoid size at either 24 or 96 h when measured with bulk fluorescence (**Figure S1, S2**). These results are not consistent with previous findings that Dps causes nucleoid compaction. However, previous assays have been performed either in fixed cells (7) or in lysed cells using atomic force microscopy (20, 36). We therefore hypothesised that Dps-mediated nucleoid compaction is more subtle *in vivo* compared with the compaction measured in previous assays and that bulk fluorescence imaging lacks the resolution needed to measure differences in nucleoid sizes between *wt* and *Δdps* cells.

To measure the nucleoid size with higher resolution, we applied live-cell single-molecule super-resolution (PALM) imaging using an *E. coli* strain that expresses HUα-PAmCherry from the native promoter. HUα binds DNA without sequence specificity (37), and PAmCherry is a photo-activatable fluorescent protein that does not interfere with HUα function (38). Moreover, HUα-mCherry has been shown to highly colocalise with the nucleoid and to well define nucleoid structure (39, 40). Therefore, HUα-PAmCherry is frequently used as a non-specific nucleoid label for super-resolution imaging (38, 41–43). From PALM images of the nucleoid based on HUα-PAmCherry localisations in stationary and deep stationary phases for dozens of *wt* and *Δdps* cells (**Figure 2B**), we quantified the nucleoid occupancy (nucleoid area divided by cell area) for *wt* and *Δdps* cells (**Figure 2C**). At 24 h, these distributions are not statistically distinct, consistent with our results from bulk fluorescence microscopy (**Figure S1**). Aligned with previous reports (40, 44), we find that the nucleoid occupancy measured by SYTOX green (a small molecule) and HUα-PAmCherry is similar (**Table S5**), despite the bulky size of PAmCherry. However, single-molecule fluorescence microscopy can resolve nucleoid compaction due to Dps in deep stationary phase (96 h): at 96 h, the *wt* and *Δdps* distributions are statistically distinct. Furthermore, at 96 h, the *wt* nucleoid occupancy distribution widens considerably (**Figure 2C**): though most of the cells at 96 h have a nucleoid that is either mildly compacted or not compacted due to Dps, ∼20% of cells in the *wt* population have deeply compacted nucleoids (nucleoid occupancy < 0.4) (**Figure S3**). This heterogeneity in the nucleoid occupancy at 96 h for *wt* cells is consistent with the work of Kovalenko *et al*. who found that only ∼20% of *wt* cells have clear Dps-DNA co-crystals in Cryo-EM (23).

### Dps asymmetrically compacts the *E. coli* nucleoid

We pooled the super-resolution localisations from all *wt* and *Δdps* HUα-PAmCherry cells and plotted them as two-dimensional heatmaps within a normalised cell area (**Figure 2D**). At 96 h, HUα-PAmCherry has a similar spatial distribution along the cell long axis (*x*-axis) for *wt* and *Δdps*, but the HUα-PAmCherry distribution shows that the *wt* cell nucleoids are more compacted along the *y*-axis compared to *Δdps* cells (**Figure 2D, 2E**). This asymmetric compaction is not clearly observed at 24 h (**Figure S4**). Our data indicate that Dps is more likely to compact the nucleoid along the short axis of cells, rather than affecting the nucleoid length.

### Dps decreases nucleoid accessibility for large protein assemblies at 24 h

We reasoned that Dps-mediated nucleoid compaction could decrease nucleoid mesh size, thereby sterically restricting proteins and particles from diffusing through the nucleoid. To test the hypothesis that Dps influences the nucleoid accessibility of large protein assemblies, such as ribosomes (21-nm diameter) (45), we tracked 25-nm eGFP-labelled protein nanocages (26) in *wt* and *Δdps* cells (**Figure 3A, 3B**). This probe has been previously used in a similar context by Xiang *et al*. to determine that the *E. coli* nucleoid mesh size is greater than 25 nm in exponential phase, as these probes were not excluded from exponential-phase nucleoids (24). The nanocages do not interact specifically with any subcellular components of *E. coli* (24), allowing us to isolate effects due solely to steric effects originating from nucleoid compaction.

Heatmaps of nanocage localisations in stationary phase reveal that nanocages are primarily nucleoid-excluded: we observe low localisation probability within the nucleoid region, which is outlined in red (**Figure 3A**) and is calculated from the HUα-PAmCherry heatmap measured at 24 h (**Figure S4**). Rather, nanocages have the highest probability of localising just outside the nucleoid, toward the cell poles (yellow pixels, **Figure 3A**). Our finding that 25-nm probes are nucleoid-excluded indicates that the nucleoid mesh size decreases in stationary phase relative to exponential phase, during which 25-nm nanocages were equally distributed throughout the cell (24). This indication that the nucleoid exhibits altered structure in stationary phase is perhaps unsurprising, considering that NAP expression dramatically changes in stationary phase (12), and we find that the chromosome conformation restructures by 24 h of starvation (**Figure 6**).

From the nanocage heatmaps, we calculated the fraction of nanocages excluded from the nucleoid region. We defined the nucleoid regions based on the HUα-PAmCherry heatmaps collected for *wt* and *Δdps* cells at 24 and 96 h (**Figure S4, 2D**). Because the nucleoid size and shape differ in *wt* and *Δdps* cells, the fraction of excluded nanocages can fluctuate based on nucleoid geometry rather than actual differences in accessibility. Consequently, we normalised the excluded fraction to consider the three-dimensional geometry of the cell and nucleoid by calculating the theoretical maximum and minimum excluded fractions based on the nucleoid shapes (**Figure 3C)**. On this scale, a value of −1.0 would indicate all localisations occur within the nucleoid; zero would indicate localisations are uniformly distributed throughout the cell; and 1.0 would indicate that particles are entirely nucleoid-excluded. The average normalised excluded fraction of nanocages in *wt* cells at 24 h is 0.35 (**Figure 3C**), consistent with the qualitative observation that, though some nanocages are localised within the nucleoid, nanocages are more likely to be found in the cell cytoplasm (**Figure 3A**). In *Δdps* cells at 24 h, the average normalised excluded fraction is only 0.11 (**Figure 3C**), meaning that in cells lacking Dps, nanocages have a higher probability of localising within the nucleoid relative to *wt* cells, consistent with the higher localisation density within the nucleoid of *Δdps* cells compared to *wt* cells **(Figure 3A)**. Considering that Dps does not measurably compact the nucleoid at 24 h (**Figure 2C**), it is somewhat surprising that Dps would reduce nucleoid accessibility of large protein assemblies. This measurement suggests that increased steric hindrance in the nucleoid can occur just from high Dps expression at 24 h, rather than requiring an overall decrease in nucleoid occupancy.

In deep stationary phase, nanocages are equally likely to be excluded from *wt* and *Δdps* nucleoids (**Figure 3D**). The average relative excluded fraction for *wt* cells increases slightly between 24 and 96 h (0.35 to 0.50), and for *Δdps* cells, this value increases from 0.11 at 24 h to 0.38 at 96 h, which is not significantly different from the normalised excluded fraction for *wt* cells at 96 h (**Figure 3E**). This increased nucleoid exclusion for *Δdps* cells indicates that cells can compensate for the loss of Dps in deep stationary phase to maintain similar nucleoid mesh sizes for both *wt* and *Δdps* nucleoids, though these compensation mechanisms either are not activated or cannot fully compensate for the absence of Dps at 24 h. We note that the nucleoid accessibility measurements are population-averaged and that there is considerable cell-to-cell variability in nanocage localisation probability. Examining single-cell nanocage localisation maps, we found that while some cells have accessible nucleoids, others have very few nanocage localisations in the nucleoid region (**Figure S5**). Future work should explore whether nucleoids that are tightly compacted by Dps are also less permeable to nanocages.

### The Dps-coated nucleoid remains accessible to small DNA-binding proteins

Previously, Janissen *et al.* showed that *in vitro*, Dps-coated DNA is protected from cleavage by several restriction enzymes yet remains accessible to RNA polymerase (7). This result led to the proposed mechanism that Dps-DNA interactions form a phase-separated nucleoid that is selectively soluble for RNA polymerase but excludes other DNA-binding proteins like restriction enzymes (**Figure 1**). This mechanism differs from the exclusion of the nanocages described above, as restriction enzymes are too small to be sterically excluded from the nucleoid. Rather, this phase-separation model proposes that *E. coli* has evolved to concentrate proteins necessary for survival near the nucleoid while non-essential ones are excluded.

We tested this model in cells by localising and tracking single molecules of RNA polymerase and the restriction enzyme EcoRI fused to PAmCherry. Consistent with the selective solubility model, we found that *wt* and *Δdps* nucleoids are equally accessible to RNA polymerase at both 24 and 96 h (**Figure S6**). Note that RNA-seq transcriptome analysis from Janissen *et al.* found no significant difference in the expression of any individual genes between *wt* and *Δdps* cells; therefore, Dps also does not measurably impact the transcriptional activity of RNA polymerase (7). We observed that *in vitro*, Dps protects DNA from EcoRI digestion, and this protection occurs at concentrations near the DNA-dissociation constant of Dps, indicating that the protection originates from Dps binding to DNA (**Figure S7A, B**). Next, we expressed PAmCherry-EcoRI *in vivo* from a plasmid with an arabinose-inducible promoter. When PAmCherry-EcoRI is induced in otherwise *wt* cells with 0.025% arabinose in exponential-phase medium, we found a two-log reduction in CFUs after 96 h of cultivation relative to cells cultured without arabinose and *wt* cells without the EcoRI-expression plasmid (**Figure S7C**), indicating that PAmCherry-EcoRI maintains its cytotoxic nature.

To measure the single-molecule dynamics of this restriction enzyme in living cells, we avoided this cytotoxicity by tracking PAmCherry-EcoRI[E111Q], a mutant that cannot cut DNA but binds DNA 1000× more strongly than the native protein (46) in *wt* and *Δdps* cells at the 96-h time point (**Figure 3F**). In both *wt* and *Δdps* cells, PAmCherry-EcoRI[E111Q] has the highest localisation probability within the nucleoid (red outline in **Figure 3G**). The normalised excluded fractions of PAmCherry-EcoRI[E111Q] in *wt* and *Δdps* cells are negative (i.e., biased toward the nucleoid) and are not significantly different (**Figure 3H**), confirming that the molecules are preferentially enriched within the nucleoid with or without Dps. Furthermore, PAmCherry-EcoRI[E111Q] molecules diffuse slowly, similar to chromosomal loci (**Figures S7D**), indicating that PAmCherry-EcoRI[E111Q] molecules strongly bind DNA, as expected for this EcoRI mutant. Together, our results indicate that PAmCherry-EcoRI[E111Q] can access and bind Dps-bound DNA in *wt* cells with the same efficiency as in cells lacking Dps. In other words, we do not observe evidence for Dps-DNA interactions driving selective insolubility of EcoRI within the *E. coli* nucleoid, as was previously hypothesised (7). Finally, we confirmed that under our induction conditions, Dps does not protect against EcoRI damage *in vivo* (**Figure S7E**). We reconcile the *in vitro* experiments (**Figure S7**) (7) and these *in vivo* experiments by assuming that, in living cells, additional factors such as other NAPs, crowding and salt concentrations not currently captured by *in vitro* assays prevent Dps from protecting DNA from digestion by EcoRI.

### Chromosome dynamics strongly depend on the growth phase but not on Dps

Given the current model that Dps-DNA interactions can restructure the nucleoid into crystalline or toroidal structures (23), we hypothesised that Dps may slow the diffusion and increase the confinement of chromosomal DNA (22). To determine the impact of Dps on chromosome dynamics, we constructed four strains, each with a fluorescently labelled ParB-*parS* locus within one of four different macrodomains: the origin of replication (*ori*), terminus (*ter*), left domain or right domain (47, 48). We tracked the ParB-YGFP foci using fluorescence microscopy in *wt* and *Δdps* cells and acquired images in a 2-s time interval (**Figure 4A - C**). Most stationary phase cells have fewer than two chromosome copies per cell and therefore have only one or two ParB-YGFP foci. To ensure cells with similar DNA content were compared, we separately analysed the diffusion of foci in cells with one or two foci. Considering cells with only two ParB-YGFP foci, the average diffusion coefficients of the *ori*, *ter* and left loci in *wt* cells are similar at 24 h, centred at 5 – 6 × 10^−4^ µm^2^/s (**Figure 4D**). Only the right locus mobility differs slightly, with an average diffusion coefficient around 1 × 10^−3^ µm^2^/s. Comparing the diffusion coefficients for loci in cells with and without Dps at 24 h, although the *ori*, left, and right loci exhibited significant differences (**Figure 4D**), these differences were not reproducible for a second biological replicate (**Figure S8**).

**Figure 4.**
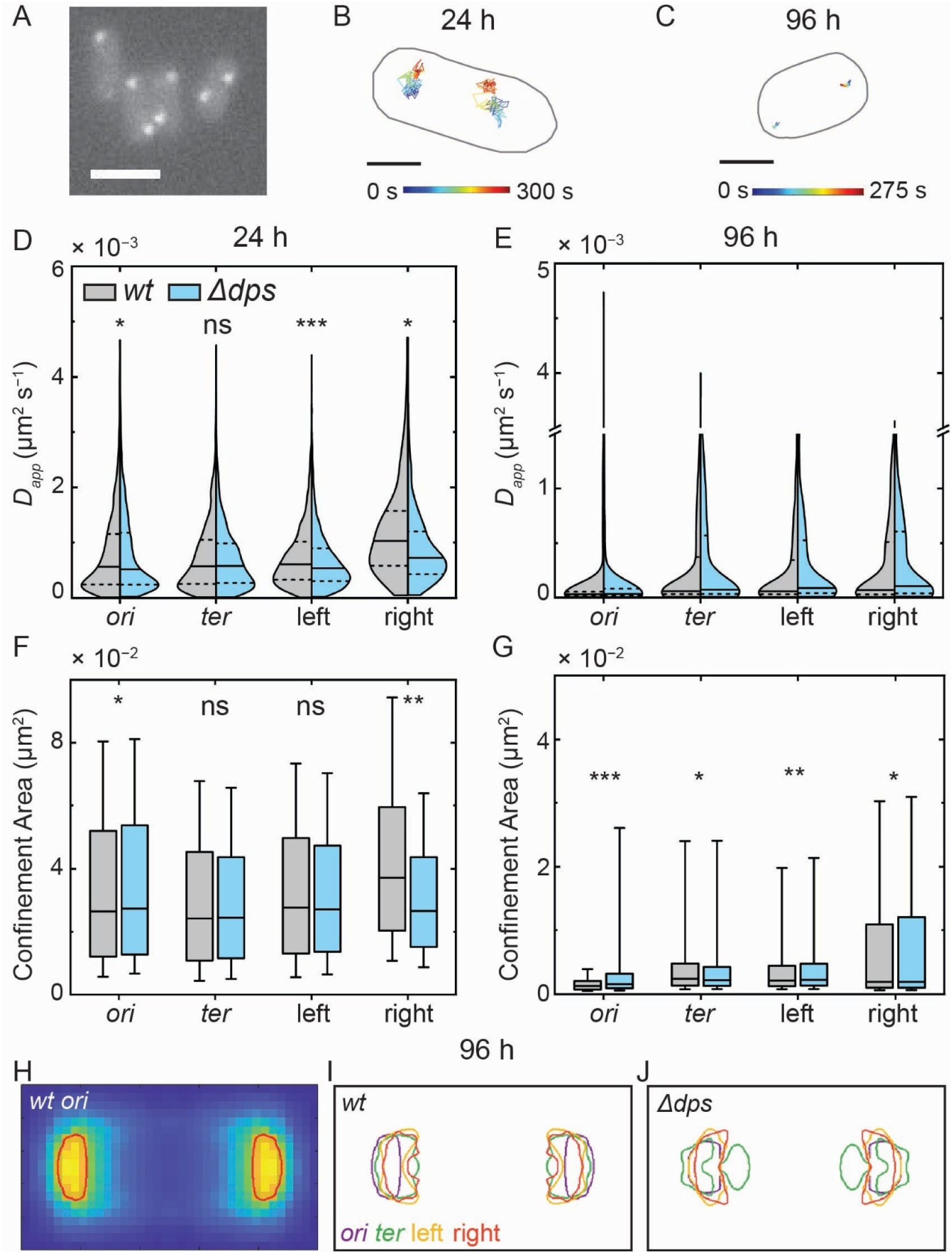
Dynamics and spatial distributions of chromosomal loci in stationary and deep stationary phase for *wt* and *Δdps* cells based on applying the ParB-*parS* labelling scheme at chromosomal loci in four different macrodomains. (A) Representative fluorescence microscopy image of ParB-YGFP labelling the ori macrodomain in *wt* cells at 24 h. Scale bar: 2 µm. Representative trajectory of a ParB-YGFP focus in *wt* cells at **(B)** 24 h and **(C)** 96 h. Scale bars: 0.5 µm. **(D)** Diffusion coefficient distributions of loci within the *ori*, *ter*, left and right macrodomains in *wt* and *Δdps* cells in stationary phase. Dashed lines mark the first and third quartiles of each distribution, and the solid line is the median. *n* ≥ 1900 ParB-YGFP foci for the *ori*, *ter* and left macrodomains and *n* ≥ 200 foci for the right macrodomain. To determine whether the diffusion coefficient distribution for each locus significantly depends on Dps, the median value of four bootstrapped subsamples was computed for each distribution; each subsample contained a quarter of the original data set. The four medians for the *wt* and *Δdps* subsampled distributions were then compared with a two-sample t-test for each locus. *: distributions determined to be significantly different (*p* < 0.05), ***: *p* < 0.001. **(E)** Same as (A), but for the 96-h time point. *n* ≥ 400 ParB-YGFP foci for each distribution. Statistical testing was not performed because > 80% of the measured diffusion coefficients are below our resolution limit. **(F)** Box and whisker plots of the confinement area of the four loci from *wt* and *Δdps* cells in stationary phase. The box length corresponds to the interquartile range, the solid line indicates the median, and the whiskers cover the 10 – 90% range of the distribution. Statistically significant differences in the confinement area were identified as in **(D)**. **(G)** Same as **(F)** but for 96 h of starvation. *: *p* < 0.05, **: *p* < 0.01, ***: *p* < 0.001. **(H)** Heatmap of the *ori* localisation density with an overlaid red outline corresponding to the top 25% of localisation probabilities, which we refer to as the “top 25% contour line”. Top 25% contour lines for each locus in *wt* (I) and *Δdps* cells **(J)** after 96 h of starvation. Heatmaps from *n* ≥ 1000 cells with two ParB-YGFP foci per cell were used to calculate the contour lines for each locus.

In deep stationary phase, the average diffusion coefficient for all loci is ∼3 – 8 × 10^−5^ µm^2^/s (**Figure 4E**). We detect a nearly ten-fold reduction in average locus diffusion coefficient between stationary and deep stationary phases. Our results are consistent with the order-of-magnitude reduction in locus diffusion coefficient previously reported by Zhu *et al*. between cells in exponential phase and cells at 48 h (41), which we also measured (**Figure S9**). In deep stationary phase, any dependence of locus diffusion on Dps is not resolvable as our localisation precision and sampling rate preclude accurate measurements of diffusion coefficients smaller than ∼ 6 × 10^−4^ µm^2^/s (**Figure S8**).

Since the 2-s imaging interval is biased to capture only the diffusion of slow molecules, we investigated if Dps leads to different chromosome dynamics on a faster timescale by tracking the loci continuously with a 40-ms frame time. However, we could not resolve any Dps-driven differences in locus diffusion (**Figure S10**) at this timescale either.

Next, we complemented diffusion coefficient analysis with measurements of the impact of Dps on the confinement area of each locus. At 24 h, the *ori* is more confined in *wt* cells than in *Δdps* cells, while the right locus is less confined in *wt* cells (**Figure 4F**). At 96 h, the average confinement area for all loci (∼ 2 × 10^−3^ µm^2^) is reduced by an order of magnitude relative to that observed at 24 h (∼ 2 – 3 × 10^−2^ µm^2^) (**Figure 4G**), consistent with the significant reduction in locus diffusivity measured at 96 h (**Figure 4E**). The *ori* is significantly more confined in *wt* cells at 96 h, while the *ter* is less confined in *wt* cells. The confinement areas of the left and right loci did not depend on Dps in a second biological replicate (**Figure S8**). Overall, chromosomal diffusion slows by an order of magnitude in deep stationary phase relative to stationary phase, and loci are ∼10× more confined. We find that Dps does not consistently affect locus confinement: Dps increases the confinement of the *ori* while having either no effect or decreasing the confinement of the other loci. This mild effect of Dps is consistent with its relatively minor impact on chromosome dynamics and accessibility.

We separately considered cells with only one ParB-YGFP focus per cell, reasoning that stressed cells are likely to have fewer chromosome copies due to stalled replication (**Figures S11, S12**). Dps increases the confinement of the left locus at 24 h, and the *ori* at 96 h, but our assay could not detect any other Dps-dependent effects in cells with only one ParB-YGFP focus (**Figures S11, S12**).

### The spatial arrangement of the chromosomal loci does not strongly depend on Dps

As Dps-DNA interactions have been previously implicated in spatially reorganising DNA (**Figure 1**), we next examined whether the spatial localisation of chromosomal loci depends on Dps. In the ParB focus heatmaps pooled from all cells, we identify where the locus is most often found by drawing a contour line around the top 25% of pixel intensities (**Figure 4H**). Comparing the most likely spatial arrangement in deep stationary phase for each locus in wt and *Δdps* cells (**Figure 4I and J**, respectively), we find that the *ori*, left and right loci have very similar spatial positioning at the cell quarter positions in *wt* and *Δdps* cells. This finding is consistent with our results at 24 h (**Figure S13**). In *wt* cells, the *ter* has a higher probability of localising in a compact region at the quarter positions than in *Δdps* cells (**Figure 4I, J**), but the overall spatial positioning of the *ter* locus is similar for both cell types. Therefore, we conclude that at the population scale, Dps does not play a major role in spatially organising the chromosome under our experimental conditions. However, it is possible that the specific chromosome architecture varies from cell-to-cell due to Dps, but that these differences average out when considering the whole population.

### Dps binds uniformly to the chromosome during stationary phases

To determine the genome-wide distribution of Dps proteins on the chromosome, we performed chromatin immunoprecipitation sequencing (ChIP-seq) using anti-Dps antibodies (34) on *wt* cells in exponential, stationary and deep stationary phases. Each sample was performed in two biological replicates (**Figure S14**), and the averaged results are shown in **Figure 5A**. ChIP-seq provides a map of protein binding throughout the chromosome. We found that for each growth phase, Dps is mostly evenly distributed across the entire genome, with the highest enrichment peaks at about two-fold above background (**Figure 5A**). Nonetheless, on the 10-kb scale, variations in Dps enrichment were highly reproducible between biological replicates (**Figure S14**). To quantify these variations, we calculated the standard deviations of Dps enrichment for 10-kb windows within each averaged dataset. The magnitude of the variations in Dps enrichment changed between growth phases, with a standard deviation of 0.22 for exponential phase, 0.16 for 24 h, and 0.31 for 96 h. Finally, although Dps enrichment is only weakly correlated between exponential and the two stationary phases, it is strongly correlated between 24 h and 96 h (**Figure 5B**). These data suggest that from exponential phase to stationary phase, the Dps enrichment pattern is shifted. Additionally, although the correlation between Dps enrichment at 24 and 96 h is quite high, the variation at 96 h is much higher, suggesting that the Dps enrichment pattern becomes more pronounced in deep stationary phase (96 h). A previous ChIP-seq study revealed stronger variations in Dps enrichment between different chromosomal regions (49). However, their data varied greatly between the two biological replicates presented, making it hard to draw strong conclusions. Overall, our results suggest that while Dps is generally uniformly distributed across the genome, small stretches of DNA (∼10 kb) exclude or enrich Dps binding in a sequence-dependent manner, and this effect is the most pronounced in deep stationary phase. It is unclear whether this variation is regulated in a function-specific manner.

**Figure 5.**
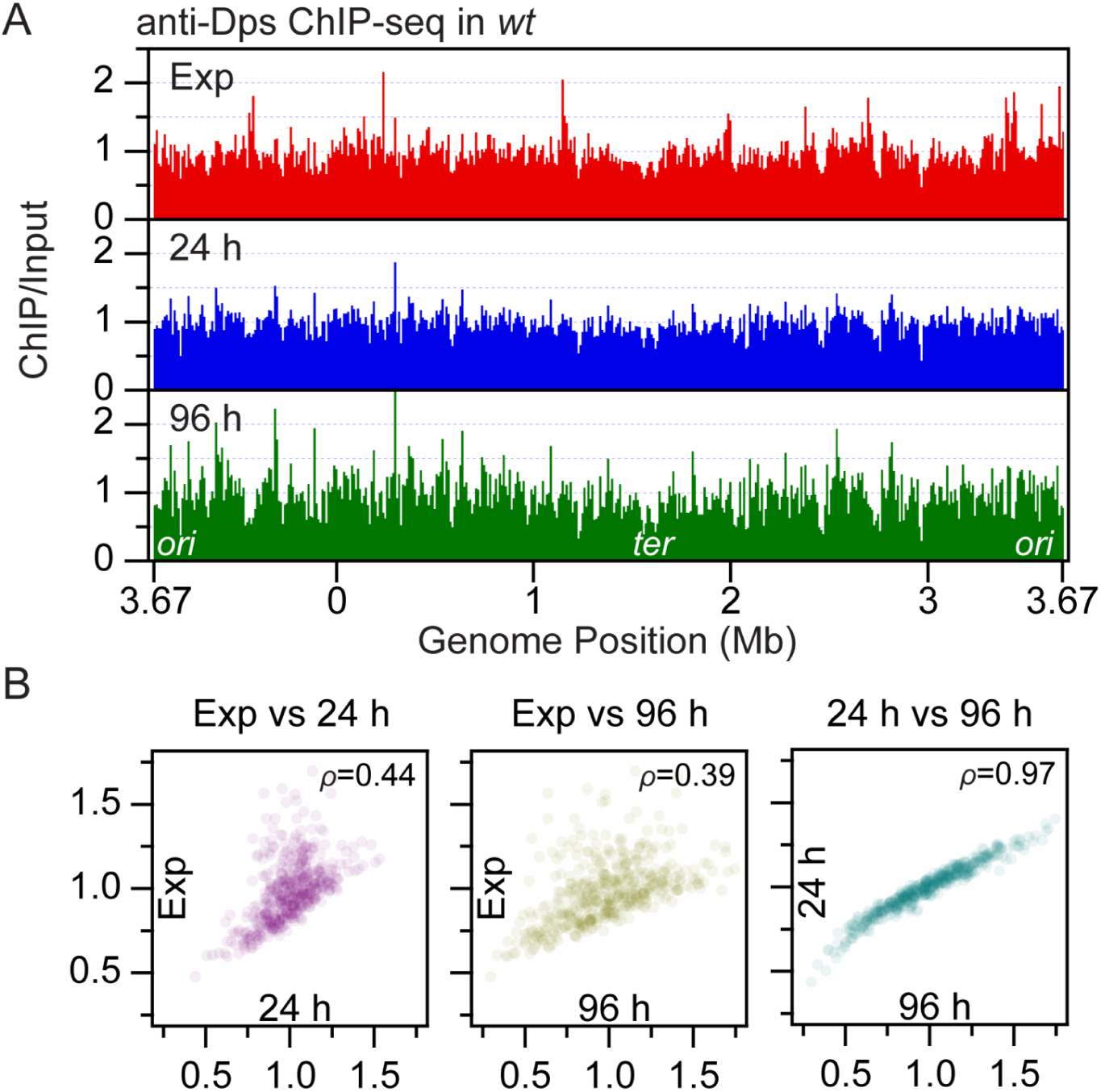
Genome-wide binding profile of Dps by ChIP-seq. **(A)** ChIP-seq of Dps in the *wt* strain for exponential phase (top panel), stationary phase (second panel) and deep stationary phase (third panel). The sequencing reads at each base pair were normalised to the total number of reads and ChIP enrichments (ChIP/input) are plotted. The genome was binned at 10-kb resolution and oriented to start at *oriC*. The average of two biological replicates is plotted here; two biological replicates for each sample are compared in **Figure S14**. **(B)** Correlation between ChIP-seq binding profiles. *ρ* is the Pearson correlation coefficient.

### The *E. coli* nucleoid is restructured in stationary phases

To understand how the three-dimensional chromosome structure responds to starvation, we performed Hi-C on *wt* cells in exponential, stationary and deep stationary phases. Hi-C combines chromosome conformation capture techniques with next-generation sequencing to provide a map of short-range and long-range DNA interactions throughout the genome. For exponential-phase *wt* cells, the Hi-C contact map has a strong primary diagonal indicating frequent short-range interactions (**Figure 6A, top left**). The Hi-C map also exhibits weaker long-range DNA contacts away from the primary diagonal, appearing as squares on the Hi-C map (dashed lines, **Figure 6A, top left**). Three major macrodomains are evident: the left replication arm, *ter* domain and right replication arm. Finally, long-range interactions nested within the macrodomains are outlined in green and correspond to smaller chromosome interaction domains (CIDs) (50) (**Figure 6A, top left**). These results are highly consistent with the work of Lioy *et al* (14). In stationary and deep stationary phases, short-range interactions progressively decrease, as evidenced by an increasingly faint primary diagonal on the Hi-C maps (**Figure 6A, top row**).

**Figure 6.**
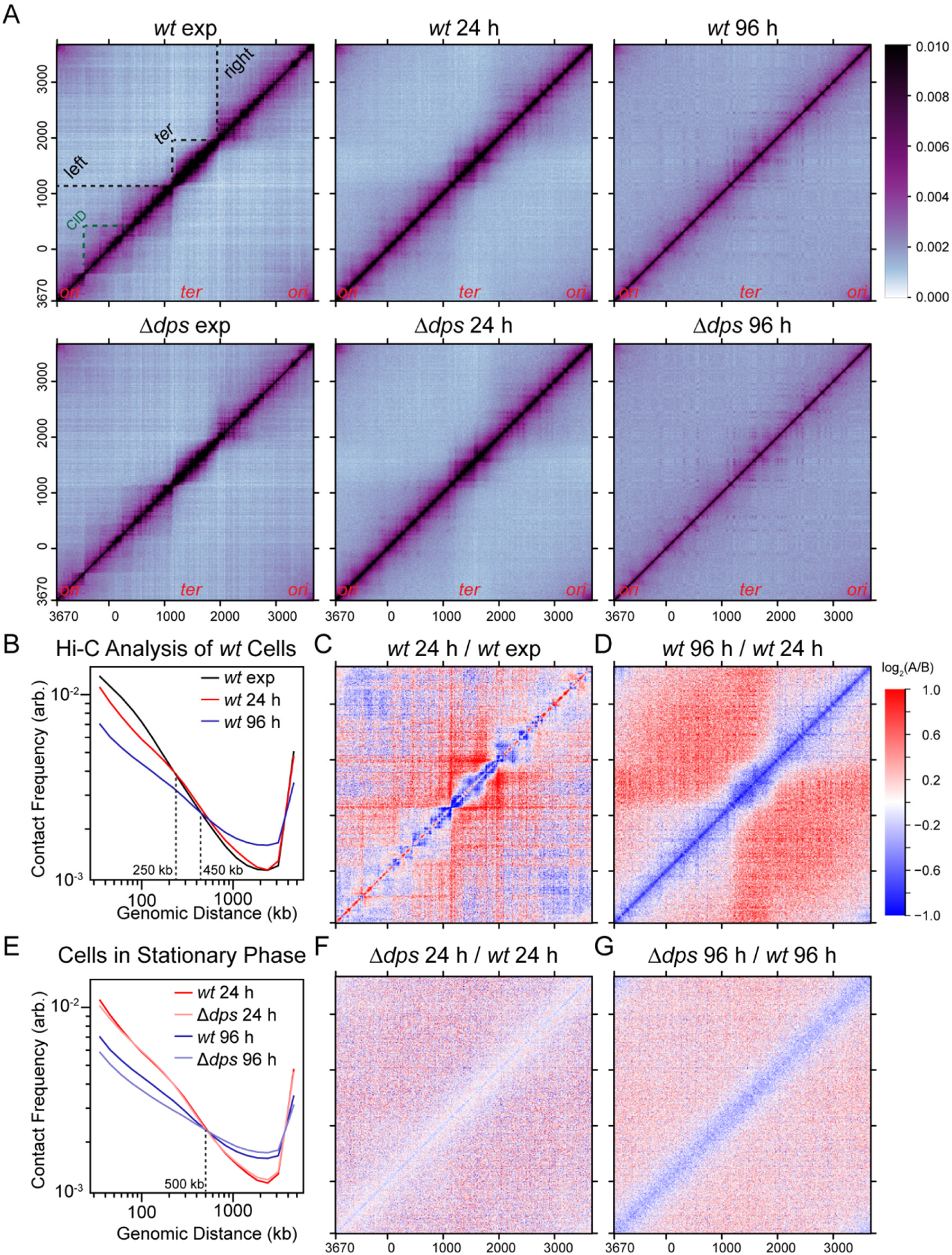
Characterisation of chromosomal interactions throughout growth phases for *wt* and *Δdps* cells. **(A)** Normalised Hi-C contact maps of *wt* and *Δdps* cells in exponential (exp), stationary (24 h) and deep stationary (96 h) phases. Black dotted lines in **(A, top left)** indicate the left, *ter* and right domains. Green dotted lines show an example of a CID. The colour bar on the right depicts Hi-C interaction scores for all contact maps. **(B)** Contact probability (Pc(s)) curves show the average contact frequency between all pairs of loci on the chromosome separated by set distances throughout the growth phases for *wt* cells. The curves were computed using data binned at 10 kb. **(C-D)** log_2_(matrix A/matrix B) ratio plots comparing different Hi-C matrices in **(A)**. **(E)** Pc(s) curves for *wt* and *Δdps* cells at the indicated time points. **(F-G)** log_2_ ratio plots comparing the indicated Hi-C matrices in **(A)**.

To quantify DNA interactions throughout the cell cycle, we analysed the average contact frequency between all pairs of loci on the chromosome separated by set distances and plotted the contact probability (Pc(s)) curves (**Figure 6B**). Compared with exponential-phase cells, stationary-phase cells exhibit reduced DNA interactions for loci separated by ∼250 kb or below; those in deep stationary phase show a further reduction in short-range interactions below ∼450 kb (**Figure 6B**). To further resolve changes in chromosome conformation as a function of growth phase, we compared the Hi-C matrices acquired at exponential and stationary phases and found a reduction in short-range interactions (**Figure 6C**, blue pixels near the primary diagonal) and notably observed the reduced presence of CIDs (**Figure 6C**, nested blue squares near the primary diagonal). Short-range interactions are further diminished between stationary phase and deep stationary phase, as is the left-*ter-*right domain structure (**Figure 6D**). These changes are also visible in the Hi-C maps (**Figure 6A, top row**). Finally, in deep stationary phase, gridded dots away from the diagonal appear on the Hi-C map, indicating that sustained interactions between multiple specific DNA regions on the chromosome cluster together (**Figure 6A, top right**). Overall, these results signify that the nucleoid is restructured in deep stationary phase, with a reduction of short-range CIDs and long-range macrodomains, but an increase in long-range specific point-to-point contacts.

### Dps facilitates short-range DNA interactions in deep stationary phase but does not restructure the nucleoid

To understand the effects of Dps on genome organisation, we compared Hi-C maps of *Δdps* cells to those of *wt* **(Figure 6A, bottom row)**. The overall genome folding patterns, such as left-*ter*-right macrodomains, CIDs and gridded dots, are very similar between *Δdps* and *wt* cells for all three growth phases **(Figure 6A)**, demonstrating that Dps is not the major driver of genome reorganisation from exponential to stationary phases. To quantify the differences in chromosome interactions between *Δdps* and *wt* cells in stationary and deep stationary phases, we calculated the Pc(s) curves and plotted the ratio of Hi-C maps **(Figure 6 E-G)**. At 24 h, the Pc(s) curves of *Δdps* and *wt* cells nearly perfectly overlap **(Figure 6E)**, and the Hi-C ratio plots show small, inconsistent differences **(Figure 6F)**. However, in deep stationary phase, *Δdps* cells exhibit reduced short-range DNA interactions between loci separated by < 500 kb compared with *wt* as shown in the Pc(s) plot **(Figure 6E)** and the blue pixels near the primary diagonal on the Hi-C ratio plot **(Figure 6G)**. These results indicate that Dps promotes local, short-range DNA interactions in deep stationary phase. This finding is consistent with the results of our super-resolution imaging, demonstrating that Dps does not significantly compact the nucleoid of live cells until deep stationary phase **(Figure 2C)**. Moreover, the finding that Dps is not the major driver of chromosome reorganisation in stationary phases is consistent with our findings that the spatial arrangement of chromosomal loci does not depend on Dps **(Figure 4I, J)**. We note that these Hi-C measurements consider the whole population and that individual cells may have chromosome organisation strongly influenced by Dps, which is averaged out at the population scale.

## DISCUSSION

In this study, we tested the current model for Dps-mediated nucleoid organisation and compaction under starvation stress (**Figure 1**) in live cells. Consistent with previous work demonstrating the protective effect of Dps (4, 11), we found that Dps increases the optical density of stationary-phase cultures relative to cells lacking Dps (**Figure 2**). We found that Dps increases nucleoid compaction in deep stationary phase using live-cell super-resolution imaging, but we also note that Dps-mediated nucleoid compaction is heterogeneous: only a portion of cells exhibit highly compact nucleoids (**Figures 2, S3**). Previous work has commented on this heterogeneity (23), but it has been under-appreciated in the field’s understanding of starved-cell nucleoid morphology. Future work should explore whether the population of cells with highly compacted nucleoids exhibits greater resistance to other stressors, such as antimicrobial agents. We further observed that Dps primarily compacts the nucleoid along the short axis of the cell (**Figure 1**). The current bottlebrush model of the nucleoid proposes that there are numerous plectonemic loops of DNA branching off from a densely packed core (51, 52). Plectonemic DNA is readily condensed by Dps *in vitro* (19), and in a 2D projection, Dps condensation of the loops could appear to compact the nucleoid to a greater extent along the short axis.

We found that Dps enhances the size-based exclusion of 25-nm nanocages from the nucleoid after 24 h of incubation (**Figure 3**), and we hypothesise that Dps binding to DNA decreases the nucleoid mesh size, leading to this greater exclusion of nanocages at 24 h. However, we expect that by 96 h, either other NAPs compensate for the absence of Dps in *Δdps* cells, decreasing the nucleoid mesh size, or the nucleoid’s structural properties similarly evolve in both *wt* and *Δdps* as transcription decreases. Therefore, *wt* and *Δdps* nucleoids equally exclude nanocages at 96 h. We found that Dps binding to DNA protects DNA from cleavage by the small restriction enzyme, EcoRI, *in vitro* (**Figure S7**), but Dps is not protective against EcoRI *in vivo* (**Figure S7**), and EcoRI is not excluded from either *wt* or *Δdps* nucleoids (**Figure 3**). Therefore, we have not observed *in vivo* evidence for the selective solubility model proposed by Janissen *et al.* (7) that predicted restriction enzymes would be insoluble in Dps-condensed nucleoids (**Figure 1**). Rather, we propose that, although Dps readily condenses DNA *in vitro* (17–19, 53, 54), in living cells, interactions with other NAPs, additional crowding factors and/or altered salt concentrations prevent Dps-DNA interactions from forming a condensate with selective solubility.

We further explored the impact of Dps on chromosome dynamics and spatial organisation by tracking genomic loci in four different macrodomains. We found that, although the growth phase strongly influences chromosome dynamics, Dps does not have a consistent impact on chromosome dynamics or spatial arrangement (**Figure 4**). Using ChIP-seq, we showed that Dps is relatively evenly dispersed throughout the chromosome at 24 and 96 h with mild sequence-dependent fluctuations in density (**Figure 5**). Despite being bound throughout the chromosome, we determined that Dps has a relatively small influence on the population-wide chromosome organisation *via* Hi-C. While chromosome interactions are dramatically altered throughout the growth phases, Dps has only a small impact on short-range DNA interactions (**Figure 6**). Together, these results indicate that, for the majority of cells, Dps does not dramatically restructure the nucleoid in stationary phase, contrary to what has been previously proposed (22). As late-stationary-phase cells have ∼180,000 Dps molecules per cell (12), it is surprising that the effect of Dps on chromosome dynamics and organisation is so modest. This subtle effect suggests that the primary role of Dps is to protect DNA through binding DNA (4) and maintaining iron homeostasis (9, 10), rather than organising the nucleoid.

Based on these results, we propose an updated model for Dps-dependent changes to nucleoid compaction, access, dynamics and organisation in response to starvation (**Figure 7**). In this model: *(a)* Dps is highly expressed at both 24 and 96 h and binds DNA, increasing nucleoid compaction at 96 h; *(b)* the nucleoid remains accessible to DNA-binding proteins for both cell types; *(c)* large protein nanocages are preferentially excluded from the *wt* nucleoid at 24 h, but by 96 h, both *wt* and *Δdps* nucleoids equally exclude nanocages; *(d)* the *wt* and *Δdps* cell chromosomes have similar organisation. It is still possible for Dps itself to exhibit a crystalline-like structure (note the regular arrangement of yellow hexagons, **Figure 7**), but perhaps Dps does not drive the underlying DNA into a crystal. Instead, Dps binds and protects DNA while only minimally impacting chromosome access, dynamics and organisation.

**Figure 7.**
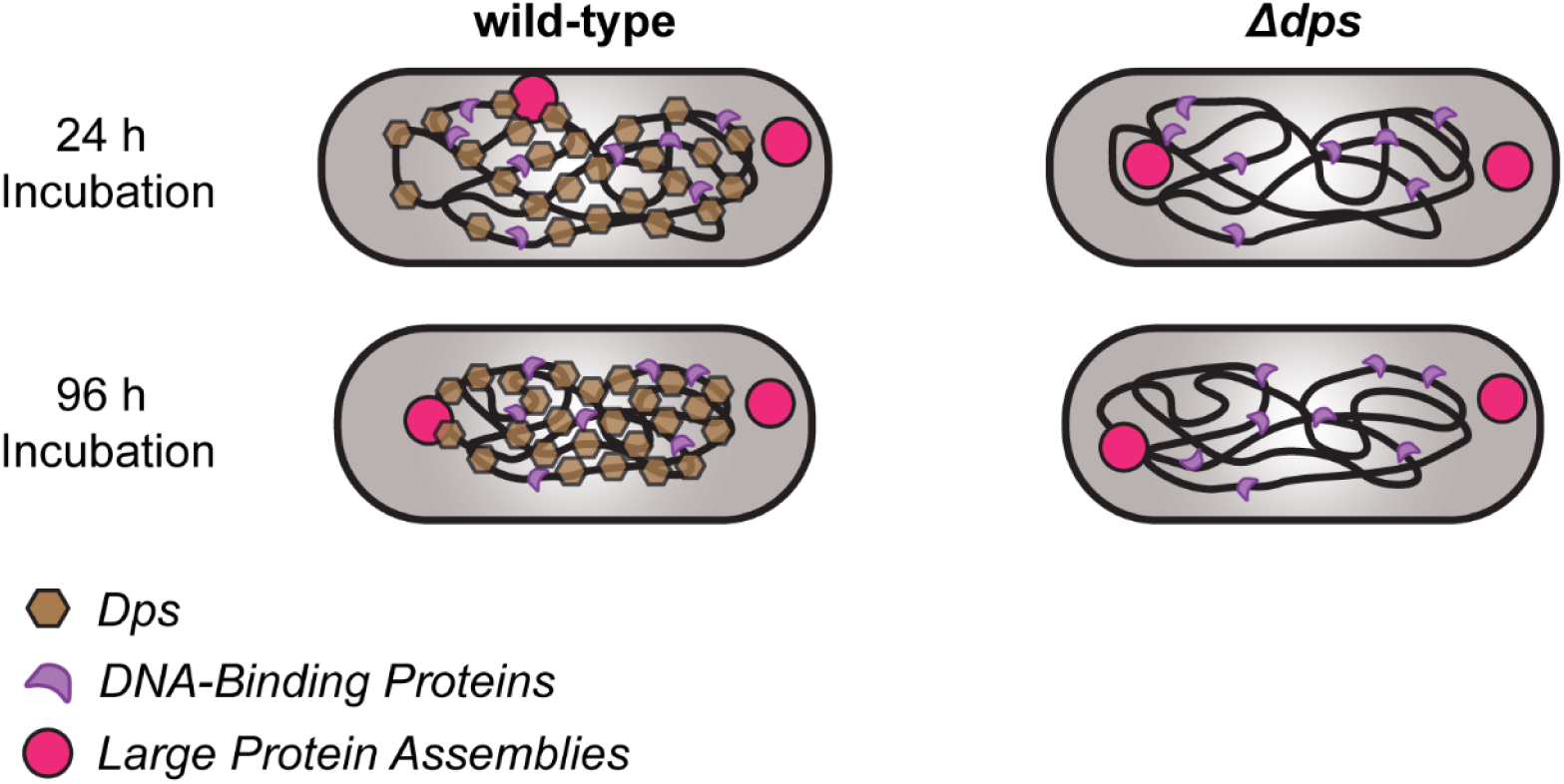
Updated model for Dps-dependent nucleoid organisation in stationary phase *E. coli* cells.

## ACKNOWLEDGEMENTS

The nanocage plasmids were a generous gift from Christine Jacobs Wagner. We thank Xheni Karaboja for technical assistance, Tom Bernhardt and Rodrigo Reyes-Lamothe for plasmids, strains and protocols, and Indiana University Center for Genomics and Bioinformatics high-throughput sequencing. The HUα-PAmCherry and RNAP-PAmCherry strains were generous gifts from Xiaowei Zhuang and Jie Xiao, respectively. We also thank Squire Booker for the pBad42 plasmid, Anthony Vecchiarelli for the PAmCherry plasmid, and Paul Modrich for the EcoRI plasmid.

## AUTHOR CONTRIBUTIONS

L.A.M. and L.E.W. conducted the investigation, developed methodology and wrote the manuscript. X.D. conducted the investigation, developed methodology and wrote analysis software. Z.R., D.E.H.F., I.D., L.L., J.J.D.S., I.W. and G.G.H. conducted the investigation and developed methodology. E.A.A. provided formal analysis of ChIP-seq data. A.S.M. supervised the study. X.W. and J.S.B. developed methodology, supervised the study and wrote the manuscript. All authors edited the manuscript.

## SUPPLEMENTARY DATA

Supplementary Data are available at NAR online.

## CONFLICT OF INTEREST

The authors declare no conflict of interest.

## FUNDING

Support for this work comes primarily from National Institutes of Health R01GM143182 (J.S.B., X.W. and A.M.). Additional support came from National Institutes of Health R01GM141242 (X.W.), R01AI172822 (X.W.), R01GM144731 (J.S.B.), and K12GM111725 (L.A.M.). Funding was also provided by the National Science Foundation MODULUS DMS-2031180 (E.A.A. and A.M.). This research is a contribution of the GEMS Biology Integration Institute, funded by the National Science Foundation DBI Biology Integration Institutes Program, Award #2022049 (X.W.).

## DATA AVAILABILITY

Hi-C, ChIP-seq and WGS data were deposited to the NCBI Gene Expression Omnibus (accession no. GSE293552; reviewer token ylynsaysfduvzmx). Single-molecule and single-particle tracking data, along with data analysis tools, were deposited to <u>Zenodo</u>, DOI: 10.5281/zenodo.15527943.

## Notes

### Competing Interest Statement

The authors have declared no competing interest.

### Summary of Updates

We have modified the manuscript with text and figure edits for clarity and new analysis (Figures 4, S5, S8, S10, and S11; Table S5).

