## Supplemental Information for "Dps binds and protects DNA in starved *Escherichia coli* with minimal effect on chromosome accessibility, dynamics and organisation"

### SUPPLEMENTAL METHODS

#### Bulk Fluorescence Microscopy of the Stained Nucleoid

The cell nucleoids were imaged on an Olympus IX-71 microscope using a CoolLED light source and a 1.40 NA 100× oil-immersion phase-contrast objective (Olympus). Cells were grown overnight at 30 °C in HDA medium and were diluted 1:100 into 50 mL of fresh HDA medium, then incubated at 30 °C. For exponential-phase imaging, samples were removed at $OD_{600}$ = 0.3. Samples were also removed at 24 and 96 h for stationary and deep-stationary-phase imaging, respectively. Per strain, three biological replicates were grown. For each replicate, 10 mL of the cell culture was extracted from the culture flask at the appropriate time and then centrifuged at 6600 × *g* for 7 min. The supernatant was collected and filtered twice with 0.22 μm filters to prepare spent medium. For exponential-phase nucleoid staining, 1 mL of exponential-phase culture ($OD_{600}$ = 0.30) was removed and stained with SYTOX green (Thermo Fisher), added to a final concentration of 500 nM. SYTOX green was chosen as the nucleoid stain as it has been previously assessed as a non-perturbative live-cell DNA stain in *E. coli* (1). The SYTOX green and cell solution was returned to the incubator and allowed to incubate for 20 min. Afterward, cells were centrifuged at 5750 × *g* for 2.5 min and washed twice with pre-warmed spent medium to remove any unbound dye. Finally, cells were resuspended in 500 μL of spent medium. For 24 and 96 h cultures, 100 μL of cell culture was added to 900 μL of spent medium and stained with SYTOX green solution added to a final concentration of 10 μM. The SYTOX green and cell mixture was then incubated for 45 min. Increasing the SYTOX green solution to 10 μM for stationary-phase cells enabled similar signal-to-noise ratios throughout the growth phases. For imaging, 2.5 μL of the stained cell solution was pipetted on 2% agarose pads made from spent medium and sandwiched between coverslips. Phase-contrast and fluorescence images were captured on a 512 × 512 EMCCD camera (Photometrics Evolve) operating continuously with a 100 ms exposure time. For imaging DNA stained with SYTOX green, the white-light illuminator was used with a 488-nm filter cube set containing an excitation filter, a dichroic mirror and an emission filter (all from Chroma). The sample was alternately illuminated with white light and 488-nm light to capture both phase-contrast and fluorescence images.

### Calculation of Nucleoid Occupancy from Images of Cells with Nucleic Acid Staining

To determine the area of the cell occupied by the nucleoid, the cell boundary was determined from the phase-contrast image by Omnipose (2) (**Figure S2A**), and the nucleoid size was measured from the SYTOX green fluorescence channel in a two-stage process. First, the intensity distribution of the second layer of pixels from the cell boundary, which is representative of the distribution of pixel intensities in the cytoplasm, was fit to a two-component Gaussian mixture model (GMM): a lower-intensity component from the cell autofluorescence background and a higher-intensity component from the weak fluorescence intensity from free (non-rigidified) SYTOX green dye in the cytoplasm (**Figure S2C, G**). A first threshold for the cytoplasmic background was set as the mean fluorescence intensity plus the standard deviation of this second component. This first threshold value was subtracted from the cell image, and all negative pixel intensity values were set to zero (**Figure S2D**).

Second, the background-subtracted intensity distribution from the boundary pixels of the remaining region was fit to a two-component GMM (**Figure S2E, H**). The lower-intensity component came from additional cytoplasmic background that was not fully accounted for in the background subtraction, and the high-intensity component gives the intensity distribution of pixels at the nucleoid periphery. The mean fluorescence intensity plus the standard deviation of this second component was therefore set as the final threshold value for assigning the nucleoid (**Figure S2F**). All pixels with intensities greater than this threshold value were considered to be within the nucleoid. The nucleoid area is the number of pixels above the second threshold (i.e., the pixels in **Figure S2F**), the cell area is the number of pixels within the Omnipose mask (i.e., the pixels in **Figure S2A**) and the nucleoid occupancy is the ratio between the nucleoid area and the cell area.

### Assessment of Colony Forming Units (CFUs)

To assess the number of CFUs/mL for a strain expressing EcoRI from the arabinose-inducible promoter in plasmids pLAM014 and pLAM015, single colonies were first obtained from LB-agar plates supplemented with 100 µg/mL spectinomycin and 0.2% glucose, which was added to suppress EcoRI expression in the plated colonies. Overnight cultures were grown in HDA medium with spectinomycin. Saturated overnight cultures were sub-cultured into 25 – 50 mL of fresh HDA medium containing 100 μg/mL spectinomycin and 0.025% arabinose to an $OD_{600}$ of 0.03 – 0.05. After 96 h of incubation, cells were diluted in fresh HDA medium with 100 μg/mL spectinomycin. 2 – 5 μL spot dilutions were plated on fresh LB-agar plates with spectinomycin and glucose. To assess the number of CFUs/mL for the *wt* strain W3110, the same procedure was repeated without antibiotics or arabinose.

***In vitro* EcoRI Protection Assay**

To assess whether Dps can protect DNA from cleavage by EcoRI *in vitro*, a 200-bp DNA fragment containing a centrally-located EcoRI cut site was used. Two 100-bp fragments were produced when the fragment was cleaved by EcoRI. 200 ng of DNA was combined with 0 – 2 μM of Dps (final concentration) in a buffer containing 5% PEG 8000 MW, 50 mM HEPES-KOH, 100 mM KCl and 4 mM $MgCl_2$. This mixture was incubated at 30 °C for 20 min to allow Dps and DNA to complex. Next, EcoRI was added to a final concentration of 0.05 Units/μL, and the mixture was incubated at 30 °C for 30 min, then at 65°C for an additional 30 min to inactivate EcoRI. Finally, to dissociate the Dps-DNA complexes, heparin was added to a final concentration of 8.7 μM, and the mixture was incubated at 30°C for 30 min. The DNA products were stained with Biotium GelGreen, run on a 1.5% agarose gel at 70 V for 2.5 h and imaged with a BioRad GelDoc EZ Imager.

**Plasmid Construction**

All restriction digestions were performed with restriction enzymes purchased from New England Biolabs.

*pLAM003* [*pBad42 I3-01-sfgfp, spec*] was generated by isothermal assembly of two column-purified DNA fragments 1) pNC (3) amplified with primers LM022 and LM024 to give *I3-01-sfgfp,* which is the labeled nanocage monomer; 2) pBad42 (4) linearised by EcoRI and SalI digestion and treated for 30 min with antarctic phosphatase. The construct was verified by whole-plasmid sequencing.

*pLAM012* [*pBad42, ecoRIR, spec*] was generated by isothermal assembly of two column-purified DNA fragments: 1) pSCC2 (5) amplified with primers LM045 and LM046; 2) pBad42 (4) linearised by EcoRI and SalI digestion and treated for 30 min with antarctic phosphatase. The construct was verified by whole-plasmid sequencing.

*pLAM014* [*pBad42 RBS ecoRIR, spec*] was generated by isothermal assembly of two column-purified DNA fragments: 1) pLAM012 amplified with primers LM050 and LM051 and treated for 1 h with DpnI, followed by amplification with primers LM052 and LM051 to form a ribosome binding site (*RBS)* 7 bp upstream of *ecoRIR*; 2) pBad42 (4) linearised by EcoRI and SalI digestion and treated for 30 min with antarctic phosphatase. The construct was verified by whole-plasmid sequencing.

*pLAM015* [*pBad42 RBS PAmCherry-ecoRIR, spec*] was generated by isothermal assembly of three column-purified DNA fragments: 1) pYH78 (6) amplified with primers LM053 and LM054, followed by amplification with primers LM052 and LM055 to give *RBS-PAmCherry* with the flexible linker *GSGSGSGSGSS;* 2) pLAM012 amplified with primers LM050 and LM051 and treated for 1 h with DpnI, followed by LM056 and LM051 to give *GSGSGSGSGSS-ecoRIR*; 3) pBad42 (4) linearised by EcoRI and SalI digestion and treated for 30 min with antarctic phosphatase. The construct was verified by whole plasmid sequencing.

*pLAM016* [*pBad42 RBS PAmCherry-ecoRIR[E111Q], spec*] was generated using the Q5® site-directed mutagenesis kit (New England Biolabs, E0554) following kit directions and amplifying pLAM015 with primers LM069 and LM070.

*pWX1016* [*pACYC T7T7 ygfp-ParB$^{MT1}$-parS FRT-kan-FRT, amp*] was generated by isothermal assembly of three column-purified fragments: 1) pWX294 (7) amplified by oWX2686 and oWX2687 to give *amp*, *pACYC*; 2) pWX950 (8) amplified by oWX2407 and oWX2408 to give *T7T7 ygfp-ParB$^{MT1}$-parS$^{MT1}$*; 3) pKD4 (9) amplified by oWX2688 and oWX2689 to give *FRT-kan-FRT*. The construct was sequenced using oWX2379, oWX2407 and oWX2426.

*pWX1082* [*pACYC T7T7 PdnaA ygfp-ParB$^{MT1}$-parS$^{MT1}$ FRT-kan-FRT, amp*] was generated by isothermal assembly of two column-purified fragments: 1) pWX1016 amplified by oWX2946 and oWX2844 to give *T7T7 ygfp-ParB$^{MT1}$-parS$^{MT1}$ FRT-kan-FRT, amp*; 2) *E. coli* MG1655 gDNA amplified by oWX2947 and oWX2948 to give *PdnaA*. The construct was sequenced using oWX2407, oWX2397, oWX2426 and oWX3021.

*pWX1121* [*pACYC PdnaA ygfp-ParB$^{MT1}$-parS$^{MT1}$ FRT-kan-FRT, amp*] was generated by isothermal assembly of two column-purified fragments: 1) pWX1082 amplified by oWX3052 and oWX418 to give *PdnaA ygfp-ParB$^{MT1}$-parS$^{MT1}$ FRT-kan-FRT*; 2) pWX1082 amplified by oWX3053 and oWX2071 to give *pACYC Amp*. The construct was sequenced using oWX2407, oWX2397, oWX2426 and oWX3021. This removed the *T7* promoter between *pACYC* and *PdnaA* from pWX1082.

**Strain Construction**

*AM257* [W3110, *Δdps FRT-kan-FRT*] was generated through lambda recombineering to replace the genomic *dps* with a counter-selectable *kan FRT* cassette. The *kan FRT* cassette was generated by amplifying the pKD13 plasmid (9) using primers AM31 and AM32 with 50 bp of homology upstream and downstream of the *dps* gene, allowing for site-directed recombination of the cassette into the genome. The resulting PCR product was DpnI digested, purified and transfected into electrocompetent W3110 (10) cells with the lambda Red helper plasmid pKD46 (9). Recombinants were selected on LB-kanamycin 25 μg/mL plates and incubated at 37 °C to remove the pKD46 plasmid. Colony PCR with primers AM34 and AM35 and western blotting verified removal of the *dps* gene.

*cWX2979* [W3110, *Δdps FRT-kan-FRT*] was generated by P1 transduction of W3110 (10) with P1 lysate from AM257. Transductants were selected for on LB-kanamycin 20 μg/mL plates.

*cWX2782* [W3110, *hupA::PAmCherry FRT-cam-FRT*] was generated by P1 transdruction of W3110 (10) with P1 lysate from strain hns-PAmCherry-cam (11). Transductants were selected for on LB-chloramphenicol 20 μg/mL plates.

*cWX3017* [W3110, *Δdps FRT-kan-FRT, hupA::PAmCherry FRT-cam-FRT*] was generated by P1 transdruction of cWX2979 with P1 lysate from strain hns-PAmCherry-cam(11). Transductants were selected for on LB-chloramphenicol 20 μg/mL plates.

*LAM045* [W3110, *Δdps FRT-kan-FRT, pBad42 I3-01-sfgfp, spec*] was generated by transforming chemically competent *cWX2979* with *pLAM003* by heat shock. Transformants were selected for on LB-spectinomycin 100 μg/mL plates. Colony PCR using primers LM006 and LM027 was performed to confirm plasmid presence.

*LAM056* [W3110, *pBad42 I3-01-sfgfp, spec*] was generated by transforming chemically competent W3110 (10) with *pLAM003* by heat shock. Transformants were selected for on LB-spectinomycin 100 μg/mL plates. Colony PCR using primers LM006 and LM027 was performed to confirm plasmid presence.

*LAM059* [W3110, *pBad42 RBS ecoRIR, spec*] was generated by transforming chemically competent W3110 (10) with *pLAM014* by heat shock. Transformants were selected for on LB-spectinomycin 100 μg/mL plates supplemented with 0.2% glucose to suppress EcoRI expression. Colony PCR using primers LM006 and LM027 was performed to confirm plasmid presence.

*LAM062* [W3110, *Δdps FRT-kan-FRT, pBad42 RBS ecoRIR, spec*] was generated by transforming chemically competent *cWX2979* with *pLAM014* by heat shock. Transformants were selected for on LB-spectinomycin 100 μg/mL plates supplemented with 0.2% glucose to suppress EcoRI expression. Colony PCR using primers LM006 and LM027 was performed to confirm plasmid presence.

LAM060 [W3110, *RBS PAmCherry-ecoRIR, spec*] was generated by transforming chemically competent W3110 (10) with *pLAM015* by heat shock. Transformants were selected for on LB-spectinomycin 100 μg/mL plates supplemented with 0.2% glucose to suppress PAmCherry-EcoRI expression. Colony PCR using primers LM006 and LM027 was performed to confirm plasmid presence.

*LAM061* [W3110, *RBS PAmCherry-ecoRIR[E111Q], spec*] was generated by transforming chemically competent W3110 (10) with *pLAM016* by heat shock. Transformants were selected for on LB-spectinomycin 100 μg/mL plates supplemented with 0.2% glucose to suppress PAmCherry-EcoRI[E111Q] expression. Colony PCR using primers LM006 and LM027 was performed to confirm plasmid presence.

*LAM064* [W3110, *Δdps FRT-kan-FRT, RBS PAmCherry-ecoRIR[E111Q], spec*] was generated by transforming chemically competent *cwx2979* with *pLAM016* by heat shock. Transformants were selected for on LB-spectinomycin 100 μg/mL plates supplemented with 0.2% glucose to suppress PAmCherry-EcoRI[E111Q] expression. Colony PCR using primers LM006 and LM027 was performed to confirm plasmid presence.

*cWX2606* [*TB10, ydeA(ter) PdnaA ygfp-ParB$^{MT1}$-parS$^{MT1}$ FRT-kan-FRT*] was generated through lambda recombineering. The *(ydeA(ter) PdnaA ygfp-ParB$^{MT1}$-parS$^{MT1}$ FRT-kan-FRT)* cassette was constructed by isothermal assembly of three fragments; 1) W3110 (10) gDNA amplified with oWX2765 and oWX2766 to make 500 bp with homology to the upstream region of *ydeA*; 2) pWX1121 amplified with oWX2815 and oWX2816 to give *PdnaA ygfp-ParB$^{MT1}$-parS$^{MT1}$ FRT-kan-FRT*; 3) W3110 (10) gDNA amplified with oWX2767 and oWX2768 to make 500 bp with homology to the downstream region of *ydeA*. The resulting fragment was gel-purified and electroporated into TB10 (12). Recombinants in which the cassette was integrated into ydeA were selected for on LB-kanamycin 20 μg/mL plates. Sequencing was performed using oWX2765 and oWX2768.

*cWX2610* [W3110, *ydeA(ter) PdnaA ygfp-ParB$^{MT1}$-parS$^{MT1}$ FRT-kan-FRT*] was generated by P1 transduction of W3110 (10) with P1 lysate from cWX2606. Transductants were selected for on LB-kanamycin 20 μg/mL plates.

*cWX2704* [W3110, *ydeA(ter) PdnaA ygfp-ParB$^{MT1}$-parS$^{MT1}$ FRT*] was generated by FLP-mediated site-specific excision of a *kan* cassette through transformation of pCP20 (13) to cWX2610. An *FRT* scar was left after cassette excision.

*cWX2724* [W3110, *ydeA(ter) PdnaA ygfp-ParB$^{MT1}$-parS$^{MT1}$ FRT, Δdps kan*] was generated by P1 transduction of cWX2704 with P1 lysate from AM257. Transductants were selected for on LB-kanamycin 20 μg/mL plates.

*cWX2564* [*TB10, yidA(ori) PdnaA ygfp-ParB$^{MT1}$-parS$^{MT1}$ FRT-kan-FRT*] was generated through lambda recombineering. The (*yidA(ori) PdnaA ygfp-ParB$^{MT1}$-parS$^{MT1}$ FRT-kan-FRT*) cassette was made by amplification of pWX1121 with oWX2981 and oWX2982. The resulting fragment contained *PdnaA ygfp-ParB$^{MT1}$-parS$^{MT1}$ FRT-kan-FRT* flanked on either side with 50 bp of DNA that is homologous to *yidA*. The fragment was column purified and electroporated into TB10 (12). Recombinants in which the cassette was integrated into *yidA* were selected for on LB-kanamycin 20 μg/mL. Sequencing was performed using oWX2407, oWX2397, oWX2426 and oWX3021.

*cWX2565* [W3110, *yidA(ori) PdnaA ygfp-ParB$^{MT1}$-parS$^{MT1}$ FRT-kan-FRT*] was generated by P1 transduction of W3110 (10) with P1 lysate from cWX2564. Transductants were selected for on LB-kanamycin 20 μg/mL plates.

*cWX2716* [W3110, *yidA(ori) PdnaA ygfp-ParB$^{MT1}$-parS$^{MT1}$ FRT*] was generated by FLP-mediated site-specific excision of a *kan* cassette through transformation of pCP20 (13) to cWX2565. An *FRT* scar was left after cassette excision.

*cWX2730* [W3110, *yidA(ori)PdnaA ygfp-ParB$^{MT1}$-parS$^{MT1}$ FRT, Δdps kan*] was generated by P1 transduction of cWX2716 with P1 lysate from AM257. Transductants were selected for on LB-kanamycin 20 μg/mL plates.

*cWX2612* [*TB10, yfeS(left) PdnaA ygfp-ParB$^{MT1}$-parS$^{MT1}$ FRT-kan-FRT*] was generated through lambda recombineering. The (*yfeS(left) PdnaA ygfp-ParB$^{MT1}$-parS$^{MT1}$ FRT-kan-FRT*) cassette was constructed by isothermal assembly of three fragments; 1) W3110 (10) gDNA amplified with oWX2769 and oWX2770 to make 600bp with homology to the upstream region of *yfeS*; 2) pWX1121 amplified with oWX2817 and oWX2818 to give *PdnaA ygfp-ParB$^{MT1}$-parS$^{MT1}$ FRT-kan-FRT*; 3) W3110 (10) gDNA amplified with oWX2771 and oWX2772 to make 750bp with homology to the downstream region of *yfeS*. The resulting fragment was gel-purified and electroporated into TB10 (12). Recombinants in which the cassette was integrated into *yfeS* were selected for on LB-kanamycin 20 μg/mL plates. Sequencing was performed using oWX2769, oWX2379, oWX2217 and oWX2772.

*cWX2621* [W3110, *yfes(left) PdnaA ygfp-ParB$^{MT1}$-parS$^{MT1}$ FRT-kan-FRT*] was generated by P1 transduction of W31101 with P1 lysate from cWX2612. Transductants were selected for on LB-kanamycin 20 μg/mL plates.

*cWX2705* [W3110, *yfeS(left) PdnaA ygfp-ParB$^{MT1}$-parS$^{MT1}$ FRT*] was generated by FLP-mediated site-specific excision of a *kan* cassette through transformation of pCP20 (13) to cWX2621. An *FRT* scar was left after cassette excision.

*cWX2726* [W3110, *yfeS(left) PdnaA ygfp-ParB$^{MT1}$-parS$^{MT1}$ FRT, Δdps kan*] was generated by P1 transduction of cWX2705 with P1 lysate from AM257. Transductants were selected for on LB-kanamycin 20 μg/mL plates.

*cWX2614* [*TB10, nlpE(right) PdnaA ygfp-ParB$^{MT1}$-parS$^{MT1}$ FRT-kan-FRT*] was generated through lambda recombineering. The (*nlpE(right) PdnaA ygfp-ParB$^{MT1}$-parS$^{MT1}$ FRT-kan-FRT*) cassette was constructed by isothermal assembly of three fragments; 1) W3110 (10) gDNA amplified with oWX2773 and oWX2774 to make 550bp with homology to the upstream region of *nlpE*; 2) pWX1121 amplified with oWX2819 and oWX2820 to give *PdnaA ygfp-ParB$^{MT1}$-parS$^{MT1}$ FRT-kan-FRT*; 3) W3110 (10) gDNA amplified with oWX2775 and oWX2776 to make 750bp with homology to the downstream region of *nlpE*. The resulting fragment was gel-purified and electroporated into TB10 (12). Recombinants in which the cassette was integrated into *nlpE* were selected for on LB-kanamycin 20 μg/mL plates. Sequencing was performed using oWX2773, oWX2379, oWX2217 and oWX2776.

*cWX2623* [W3110, *nlpE(right) PdnaA ygfp-ParB$^{MT1}$-parS$^{MT1}$ FRT-kan-FRT*] was generated by P1 transduction of W3110 (10) with P1 lysate from cWX2614. Transductants were selected for on LB-kanamycin 20 μg/mL plates.

*cWX2707* [W3110, *nlpE(right) PdnaA ygfp-ParB$^{MT1}$-parS$^{MT1}$ FRT*] was generated by FLP-mediated site-specific excision of a *kan* cassette through transformation of pCP20 (13) to cWX2623. An *FRT* scar was left after cassette excision.

*cWX2728* [W3110, *nlpE(right) PdnaA ygfp-ParB$^{MT1}$-parS$^{MT1}$ FRT, Δdps kan*] was generated by P1 transduction of cWX2707 with P1 lysate from AM257. Transductants were selected for on LB-kanamycin 20 μg/mL plates.

*cWX2695* [W3110, *rpoC::rpoC-PAmCherry FRT-kan-FRT*] was generated by P1 transdruction of W3110 (ATCC 27325) with P1 lysate from strain CC253 (14). Transductants were selected for on LB-kanamycin 20 μg/mL plates.

cWX2718 [W3110, *rpoC::rpoC-PAmCherry FRT no ab*] was generated by FLP-mediated site-specific excision of a *kan* cassette through transformation of pCP20 (13) to cWX2695. An *FRT* scar was left after cassette excision.

cWX2732 [W3110, *rpoC::rpoC-PAmCherry FRT no ab, Δdps FRT-kan-FRT*] was generated by P1 transdruction of cWX2718 with P1 lysate from AM257. Transductants were selected for on LB-kanamycin 20 μg/mL plates.

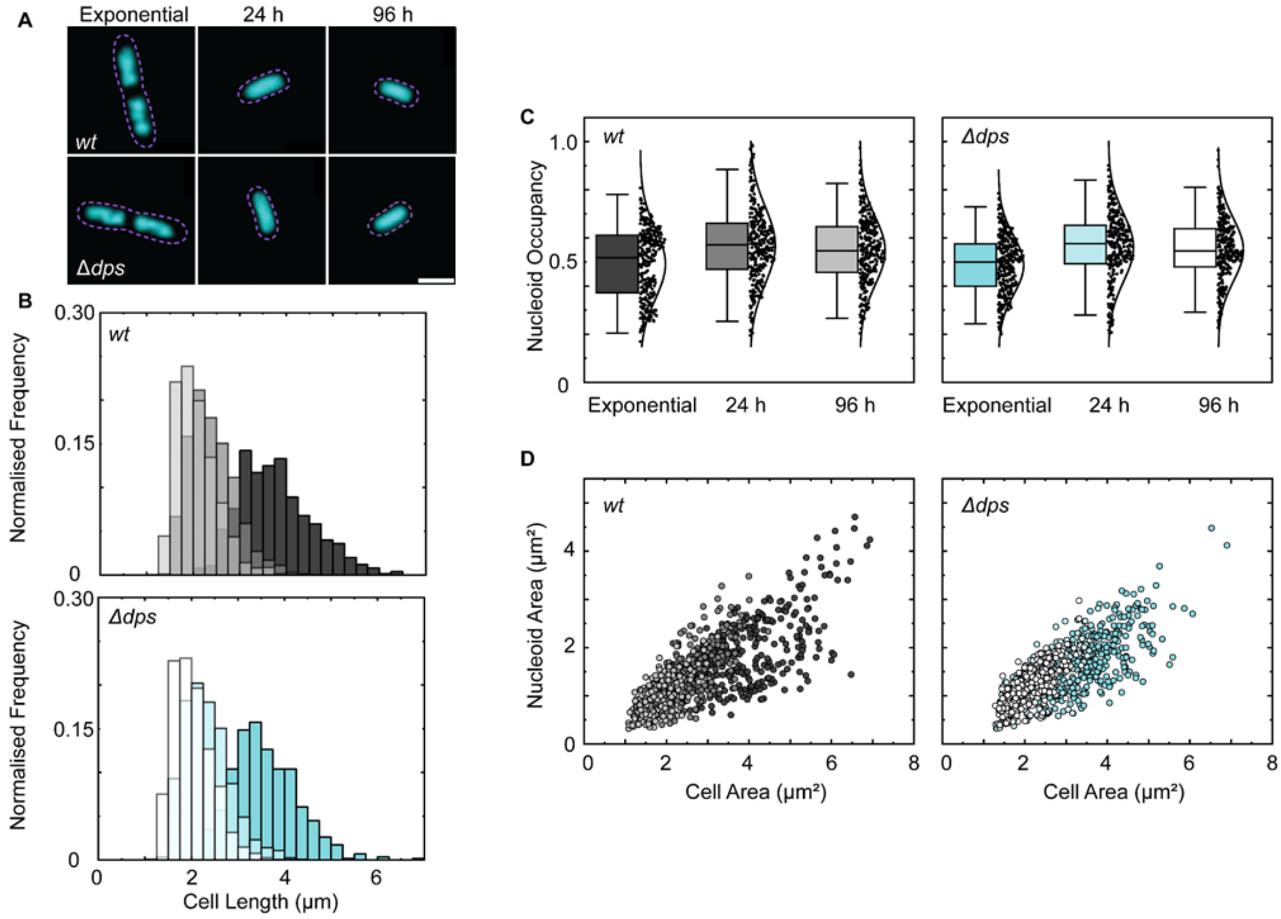

**Figure S1. Bulk fluorescence microscopy does not capture nucleoid compaction by Dps *in vivo*. (A)** Representative fluorescence images of the nucleoid stained with Sytox Green in *wt* and *Δdps* HUα-PAmCherry cells in the exponential, stationary and deep stationary phase (from left to right). The purple dotted lines are the cell outlines. Scale bar = 2 μm. **(B)** Histogram of cell length distributions of *wt* (top) and *Δdps* cells (bottom) in exponential phase (4 h) (*wt* = dark grey, *Δdps* = blue), stationary phase (24 h) (*wt* = grey, *Δdps* = light blue) and deep stationary phase (96 h) (*wt* = light grey, *Δdps* = white). Each condition contains $n$ = 1200 – 1900 cells. The mean lengths of *wt* and *Δdps* cells at 24 h were 2.40 ± 0.51 μm and 2.34 ± 0.50 μm, respectively, while at 96 h, the mean lengths were 2.07 ± 0.45 μm and 2.03 ± 0.43 μm, respectively. **(C)** Nucleoid occupancy (the nucleoid area divided by the cell area) for *wt* (left) and *Δdps* (right) HUα-PAmCherry cells. The centre line is the median and whiskers correspond to 2 standard deviations from the mean ($n$ = 300 cells per condition). The nucleoid occupancy increases from the exponential to the stationary phases for both cell types, and Dps does not affect nucleoid compaction (for each time point, *wt* and *Δdps* nucleoid occupancy distributions are not statistically distinct ($p$ > 0.05) by a two-sample t-test). **(D)** Characterisation of cell area vs. nucleoid area of *wt* (left) and *Δdps* (right) HUα-PAmCherry cells in the exponential, stationary and deep stationary phases with the same colour coding as in **B** ($n$ = 300 cells per condition). While the average nucleoid occupancy does not change between 24 and 96 h for *wt* and *Δdps* cells, the nucleoid and cell area do decrease over time. Therefore, cells maintain the same relative nucleoid-to-cell size ratio throughout the stationary phases.

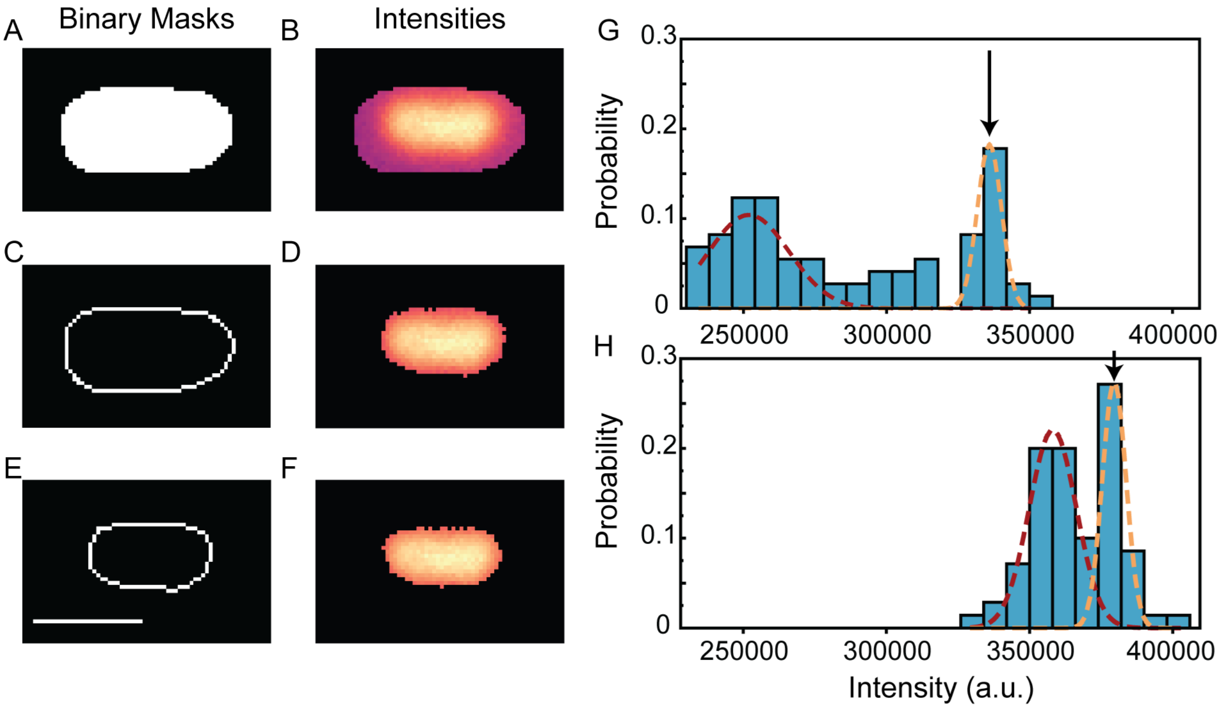

**Figure S2. Nucleoid occupancy quantification from bulk fluorescence microscopy. (A)** Binary mask of a representative stationary-phase cell from the segmentation of the phase-contrast image by Omnipose (2). **(B)** Fluorescence intensity distribution within the cell area. **(C)** The pixels of the second layer from the cell boundary, which were used to determine the distribution in panel **G**. **(D)** Fluorescence intensity distribution of the cell area remaining after eliminating all pixels with intensity less than that of the first threshold (arrow in panel **G**). **(E)** The pixels from the boundary of the area identified in panel **D**, which were used to determine the distribution in panel **H**. **(F)** The fluorescence intensity distribution of the nucleoid, which is the cell area remaining after eliminating all pixels with intensity less than the second threshold (arrow in panel **H**). Scale bar for panels **A** – **F**: 2 μm. **(G, H)** Fluorescence intensity distributions of the pixels identified in panels **C** and **E**, respectively. Each distribution is fit to a two-component Gaussian mixture model, with a dim (red) and bright (orange) component. The arrow in each panel is the threshold determined at the mean fluorescence intensity plus the standard deviation of the brighter component.

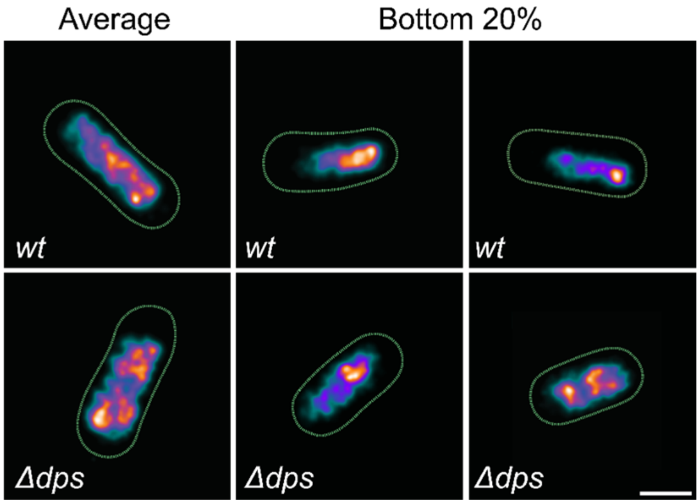

**Figure S3. Super-resolution images of representative *wt* and *Δdps* HUα-PAmCherry cells at 96 h.** Images are constructed from 1500 – 2000 localisations/cell, each with 95% confidence interval ≤ 80 nm. Colour represents the localisation density: high densities are orange and low densities are blue. Scale Bar: 1 μm. Images correspond to cells with average nucleoid occupancy and cells with nucleoid occupancy in the bottom 20% of the distribution for each cell type. As seen in **Figure 2B**, most *wt* and *Δdps* cells at 96 h have similarly sized nucleoids: the median nucleoid occupancy is 0.52 for *wt* cells and 0.56 for *Δdps*. However, ~ 20% of *wt* cells exhibit tightly compacted nucleoids: the average nucleoid occupancy in the bottom 20% of the distribution is 0.33 for *wt* cells and 0.44 for *Δdps*.

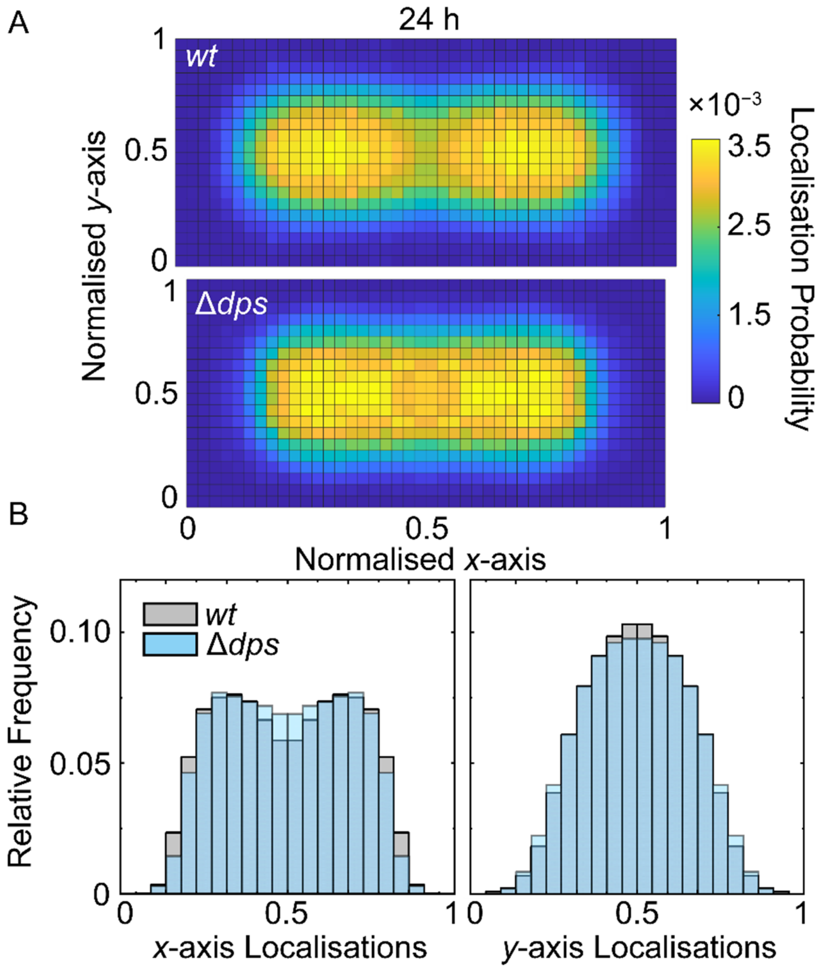

**Figure S4. HUα-PAmCherry spatial distribution at 24 h. (A)** 2D HUα-PAmCherry localisation heatmaps and **(B)** Histograms of the *x*-axis and *y*-axis localisation probabilities for *wt* and *Δdps* cells at 24 h. HUα-PAmCherry distributions for *wt* cells are slightly wider along the *x*-axis and slightly narrower along the *y*-axis compared to *Δdps* cells. Conversely, at 96 h, HUα-PAmCherry distributions for *wt* and *Δdps* cells are nearly identical along the *x*-axis, and the *wt* localisation distribution compacts only in the *y*-axis (main text **Figure 2D, E**).

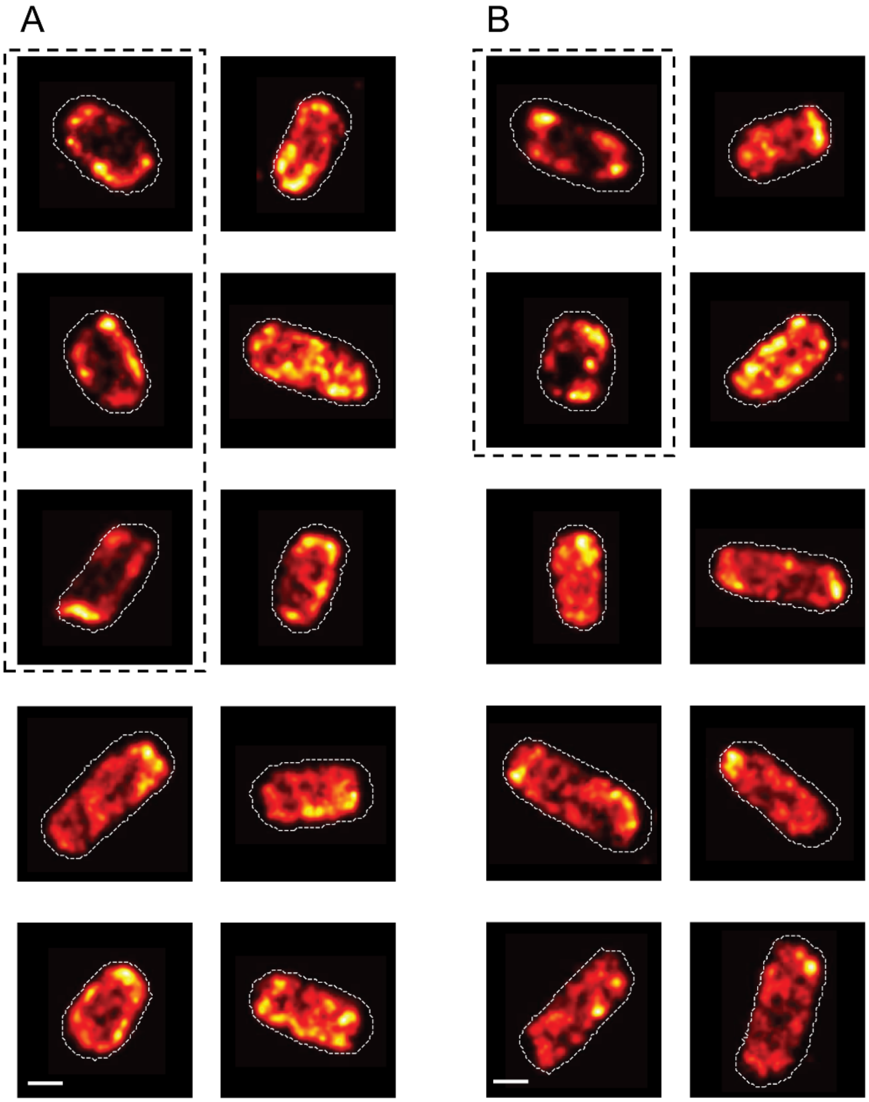

**Figure S5. Single-cell density maps of nanocage localisations at 96 h. (A)** *wt* cells exhibit two distinct nanocage localization patterns: (1) nucleoid-excluded (outlined by dashed boxes) featuring a region of low nanocage localization density at the center of the cell and (2) uniform density, suggesting relatively accessible nucleoids. **(B)** Same as A but for *Δdps* cells. Scale bar: 0.5 μm.

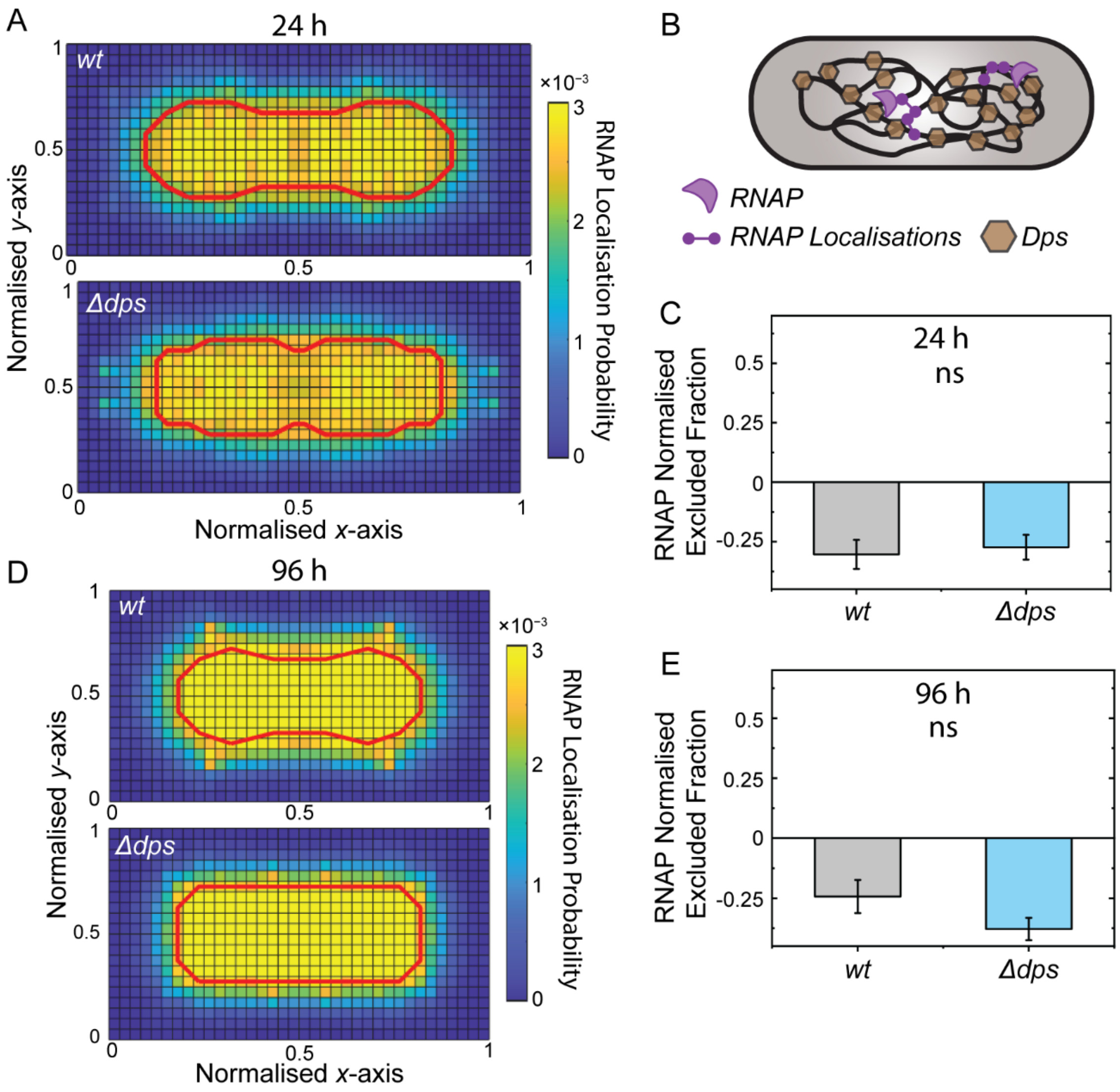

**Figure S6. Normalised excluded fraction from the nucleoid of RNA polymerase molecules in cells with and without Dps. (A)** Heatmap of drift-corrected RNA polymerase-PAmCherry localisations for *wt* and *Δdps* cells at 24 h. Drift correction was performed by tracking a nearby fluorescent bead and updating the drift every 16 – 40 s. The red outline is the top 60% contour line of localisation probabilities for HUα-PAmCherry heatmaps. **(B)** Schematic of RNA polymerase diffusing in a *wt E. coli* cell. **(C)** Normalised excluded fraction from the nucleoid of RNA polymerase molecules. The height of the bar corresponds to the mean from three randomly sampled data sets (without replacement) for $n$ ~ 10 cells each from 2 – 4 biological replicates. Error bars: standard deviations of the three sub-sampled data sets. ns: means are not significantly different ($p > 0.05$) by a two-sample t-test. **(D, E)** same as **A, C** but for cells in deep stationary phase (96 h incubation). RNA polymerase localisations have negative normalized excluded fractions at 24 and 96 h, indicating that RNA polymerase is more likely to be found in the nucleoid than the cytoplasm for both time points. The normalized excluded fractions are not significantly different for *wt* and *Δdps* cells at either time point. In other words, RNA polymerase remains nucleoid-associated throughout the stationary phase, and its nucleoid accessibility does not depend on Dps. RNA polymerase-PAmCherry data was imaged following the same single-molecule imaging protocol outlined in the main text, except with a 561-nm laser power density of 0.32 $kW/cm^2$.

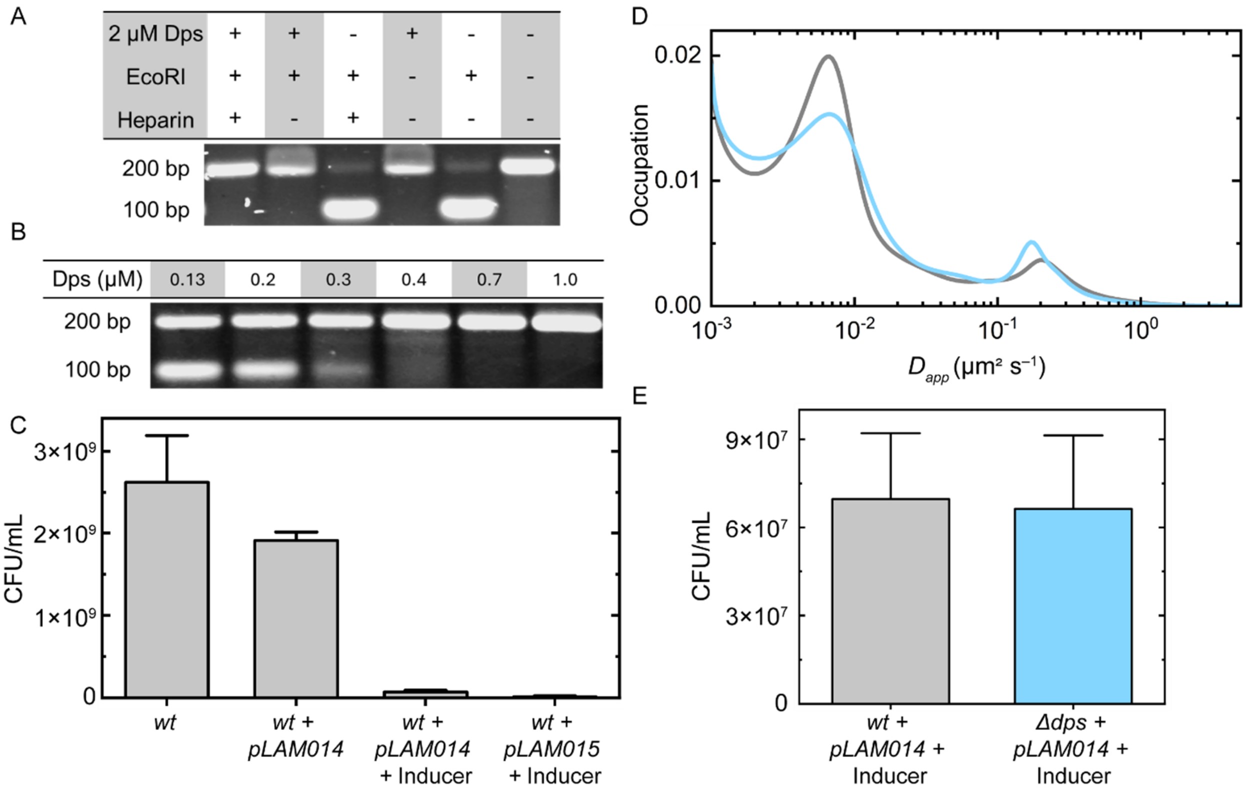

**Figure S7.** ***In vitro*, Dps prevents DNA cleavage by EcoRI. *In vivo*, EcoRI and PAmCherry-EcoRI are cytotoxic, EcoRI has slow diffusion, and Dps does not improve cell viability under EcoRI expression. (A)** Representative gel electrophoresis image of a 200-bp DNA fragment with an internal EcoRI recognition site. 200 ng of DNA was incubated with Dps (0 or 2 μM), followed by incubation with EcoRI (0 or 0.05 Units/μL). Dps-DNA complexes and DNA-EcoRI interactions were dissociated with heparin as indicated. DNA fragments pre-incubated with Dps were not cleaved by EcoRI. **(B)** Various Dps concentrations (0 – 1 μM) were incubated with 200 ng of the same DNA fragment used in **A**, followed by the addition of EcoRI (0.05 units/μL). Heparin was used to dissociate Dps-DNA complexes. Dps concentrations near or above the $K_D$ of Dps-DNA binding, which is around 0.2 – 0.6 μM (15, 16), protected DNA from cleavage by EcoRI. **(C)** CFU per volume of 96 h *wt* cells without the EcoRI-expression plasmid (*pLAM014*), with *pLAM014* but without inducer, with *pLAM014* and inducer, and with the PAmCherry-EcoRI expression plasmid (*pLAM015*) with inducer. Bar height corresponds to the mean and error bars reflect the standard deviation from 3 biological replicates. **(D)** Apparent diffusion coefficient distribution of PAmCherry-EcoRI[E111Q] measured in deep-stationary-phase *wt* and *Δdps* cells. Single-molecule trajectories were drift-corrected, and cells with fewer than 100 localisations were excluded from the analysis. Diffusion coefficient analysis was performed using the publicly available State Array algorithm (17), which uses a Bayesian framework to estimate the diffusion coefficient distribution for short single-molecule trajectories while accounting for the localisation error. $n \geq 40$ cells. The majority of trajectories have slow apparent diffusion coefficients similar to those of chromosomal loci (**Figure S10**), indicating that the majority of PAmCherry-EcoRI[E111Q] molecules are tightly bound to DNA, which is expected for this mutant (18). **(E)** Mean CFUs per volume of 96 h *wt* and *Δdps* cells with *pLAM014* and inducer. Error bars are the standard deviation from three biological replicates.

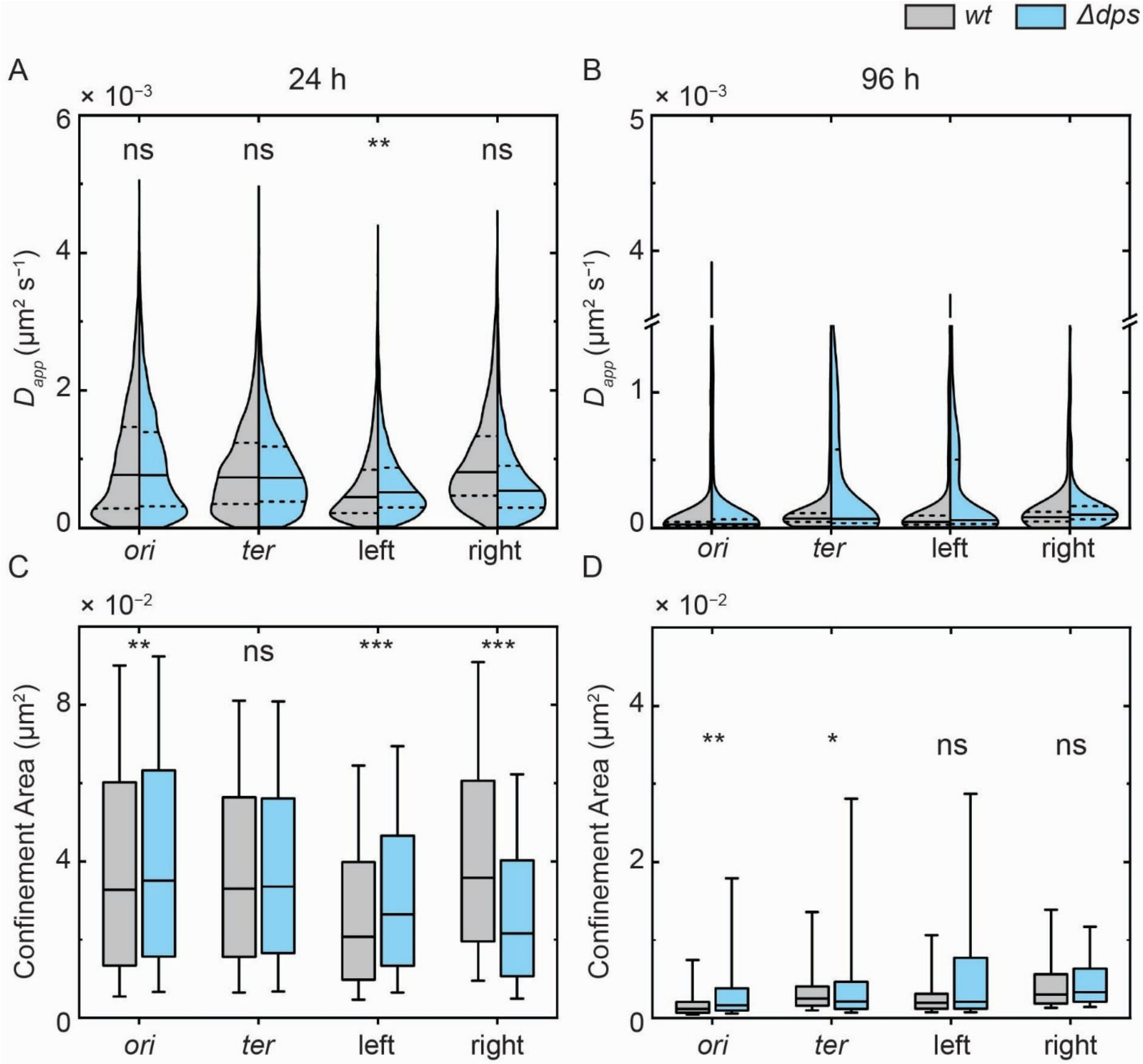

**Figure S8. A second replicate measurement of chromosomal locus dynamics at 24 and 96 h in *wt* and *Δdps* cells with 2-s time-lapse imaging. (A)** Diffusion coefficient distribution of loci within the *ori*, *ter*, left and right macrodomains after 24 h of incubation. Dashed lines mark the first and third quartiles of each distribution and solid lines are the median. $n \geq 1500$ ParB-YGFP foci for each distribution. **(B)** Same as panel **A**, but for cells incubated 96 h. $n \geq 500$ foci for the *ori, ter*, and left macrodomains and $n \geq 90$ for the right macrodomain. **(C)** Confinement area of the four loci from *wt* and *Δdps* cells at 24 h. The box length is the interquartile range, the solid line is the median and the whiskers cover the 10 – 90% range of each distribution. **(D)** Same as panel **C**, but for cells at the 96 h timepoint. To determine statistical significance, we calculated the median value of four bootstrapped subsamples (with replacement) for each distribution, with each subsample containing a quarter of the original dataset. We then compared the four medians for the *wt* and *Δdps* subsamples with a two-sample t-test for each locus. *: distributions determined to be significantly different ($p < 0.05$), **: $p < 0.01$, ***: $p < 0.001$. Statistical testing was not performed for diffusion coefficient distributions measured at 96 h because > 80% of the data in these distributions are below our localisation precision considering our localisation error (~ 0.035 μm) and sampling time (2 s). Accounting for localisation error, mean-squared displacement (*MSD*) can be calculated as $MSD = 4D\tau + 4\sigma^2$ (19), where $D$ is the diffusion coefficient, $\tau$ is the time lag and $\sigma$ is the localisation precision. To measure a reliable *MSD*, the displacement due to actual diffusion must be greater than the apparent displacement resulting from localisation error. Therefore, the slowest diffusion we can accurately quantify is: $D_{min} = \sigma^2/\tau = (0.035\ \mathrm{\mu m})^2/2\ \mathrm{s} = 6 \times 10^{-4}\ \mathrm{\mu m}^2/s$. We can, however, distinguish differences in the confinement area of the *wt* and *Δdps* distributions at 96 h because the limit of detection is ~ $\sigma^2$, which is $1.2 \times 10^{-3}\ \mathrm{\mu m}^2$.

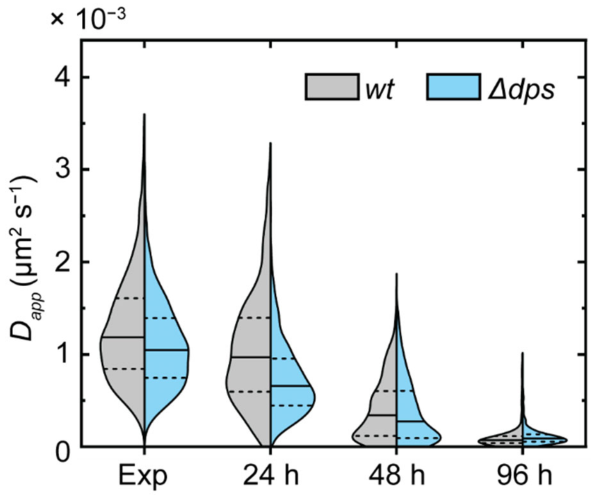

**Figure S9. Diffusion coefficient distributions of a locus in the right macrodomain in *wt* and *Δdps* cells at four different growth phases, measured through timelapse imaging with a 2-s exposure time.** Dashed lines are the first and third quartiles of each distribution. Solid lines are the medians. $n \geq 300$ cells per growth phase. Cells with different numbers of ParB-YGFP foci per cell were considered as DNA content varies throughout the growth phases. The right macrodomain has similar dynamics in the exponential phase and at 24 h, slows by 48 h, and reduces by an order of magnitude by 96 h. Dps does not strongly influence locus mobility at any of the characterised growth phases.

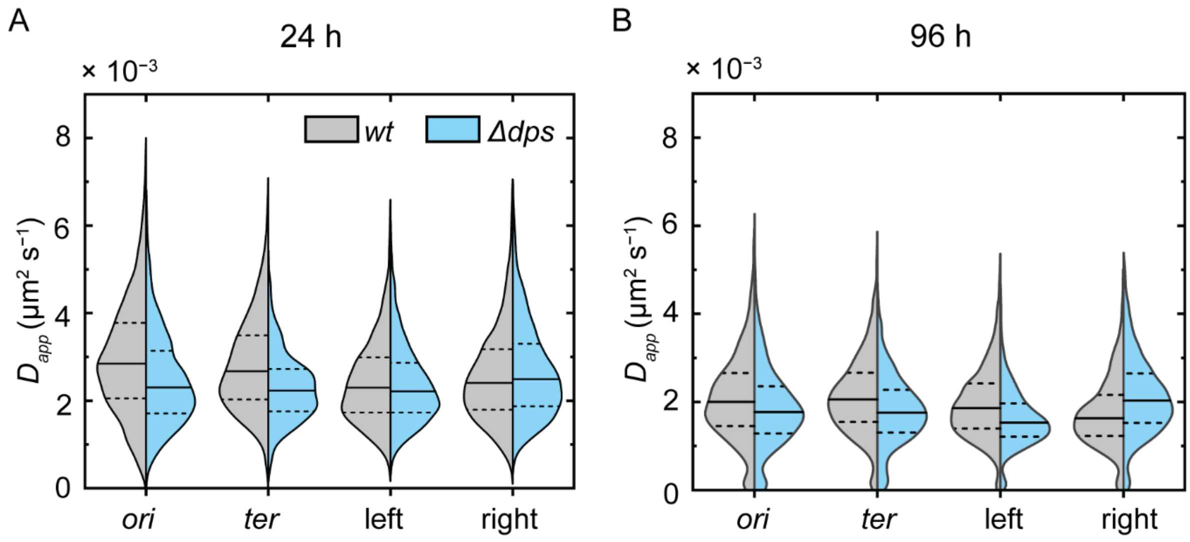

**Figure S10. Diffusion analysis from continuous imaging of chromosomal loci with a 40 ms exposure time.** Diffusion coefficient distributions of chromosomal loci within the *ori*, *ter*, left and right macrodomains in *wt* and *Δdps* cells at **(A)** 24 h and **(B)** 96 h of incubation. Dashed lines are the first and third quartiles of each distribution and the solid lines are the medians. $n \geq$ 800 ParB-YGFP foci for each macrodomain and condition. ParB-YGFP tracks from cells with two foci and that passed the 1.5 – 4.0 μm cell length filter were analysed with mean-squared displacement analysis (20). Linear fitting of the first 25% of the displacements with respect to time lag was used to calculate the diffusion coefficients. Fits with $R^2 < 0.8$ and anomalous diffusion coefficients > 1 (corresponding to superdiffusive motion not expected for chromosomal loci) were omitted from further analysis. Note that the magnitude of the apparent diffusion coefficient distributions of all loci are greater than those reported in the main text **Figure 4**. This difference in the apparent diffusion coefficient arises due to the different exposure times used: 2-s time-lapse imaging can more accurately capture slow diffusion, while 40-ms continuous imaging is sensitive to fast motion. Following the discussion in **Figure S7**, the limit of slow diffusion we can accurately capture in this assay based on our localisation error (~ 0.05 μm) and exposure time (0.04 s) is $D_{min} = (0.05\ \mu\text{m})^2/0.04\ \text{s} = 6 \times 10^{-2}\ \mu\text{m}^2/\text{s}$. As the apparent diffusion coefficient distributions for all loci at both time points are far below this cutoff, we cannot reliably distinguish differences in diffusion between 24 and 96 h or between *wt* and *Δdps* cells with a 40 ms exposure time.

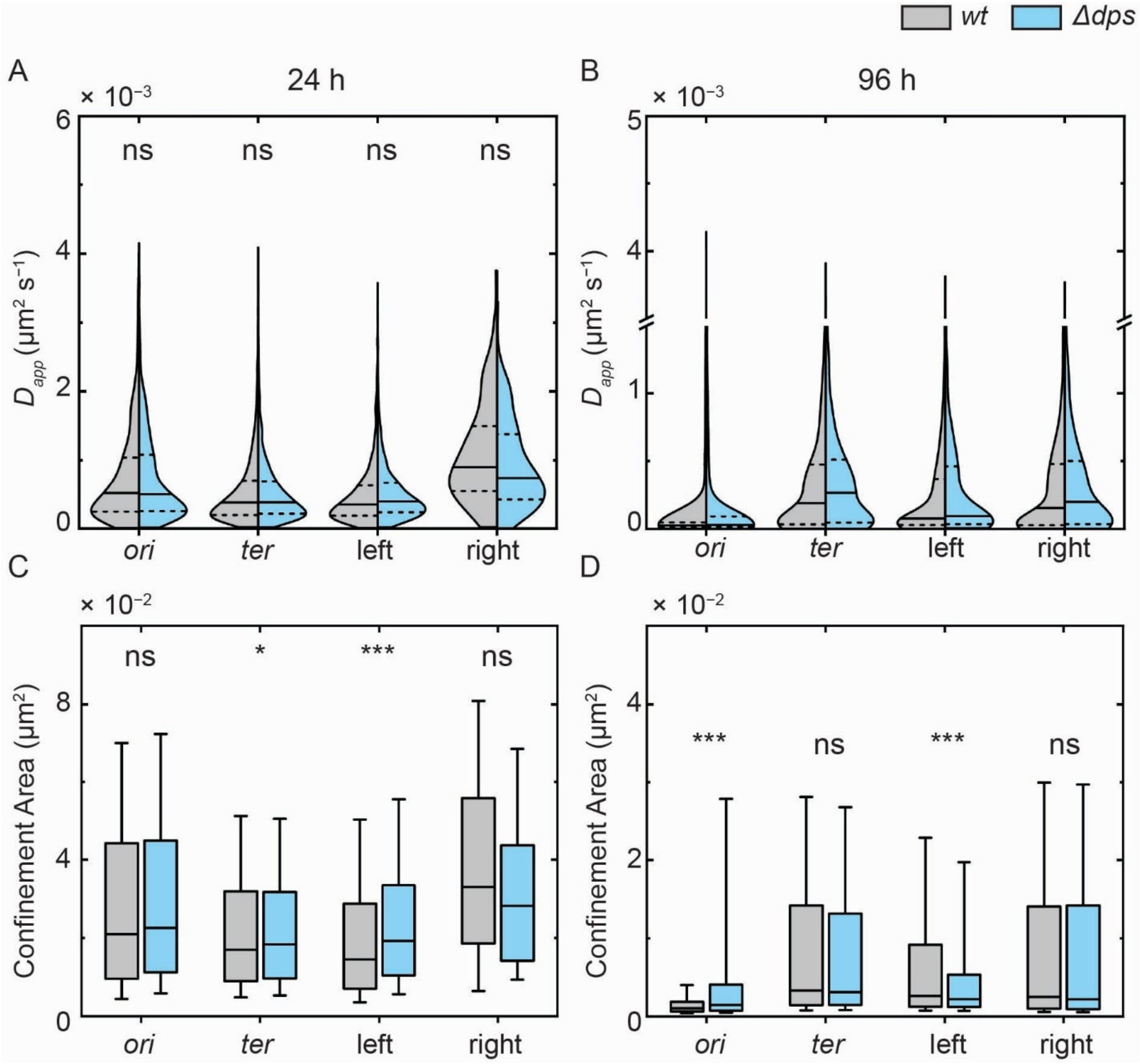

**Figure S11. Chromosomal locus dynamics at 24 and 96 h in *wt* and *Δdps* cells with only one ParB-YGFP focus with 2-s time-lapse imaging. (A)** Diffusion coefficient distribution of loci within the *ori*, *ter*, left and right macrodomains after 24 h of incubation. Dashed lines are the first and third quartiles of each distribution and solid lines are the median. $n \geq 1000$ foci for the *ori, ter*, and left macrodomains and $n \geq 90$ for the right macrodomain. **(B)** Same as panel **A**, but for cells incubated 96 h. $n \geq 750$ ParB-YGFP foci for each distribution. **(C)** Confinement area of the four loci from *wt* and *Δdps* cells at 24 h. The box length is the interquartile range, the solid line is the median and the whiskers cover the 10 – 90% range of each distribution. **(D)** Same as panel **C**, but for cells at the 96 h timepoint. Statistical analysis was performed as described above for cells with two ParB-YGFP foci. *: distributions determined to be significantly different ($p < 0.05$), **: $p < 0.01$, ***: $p < 0.001$.

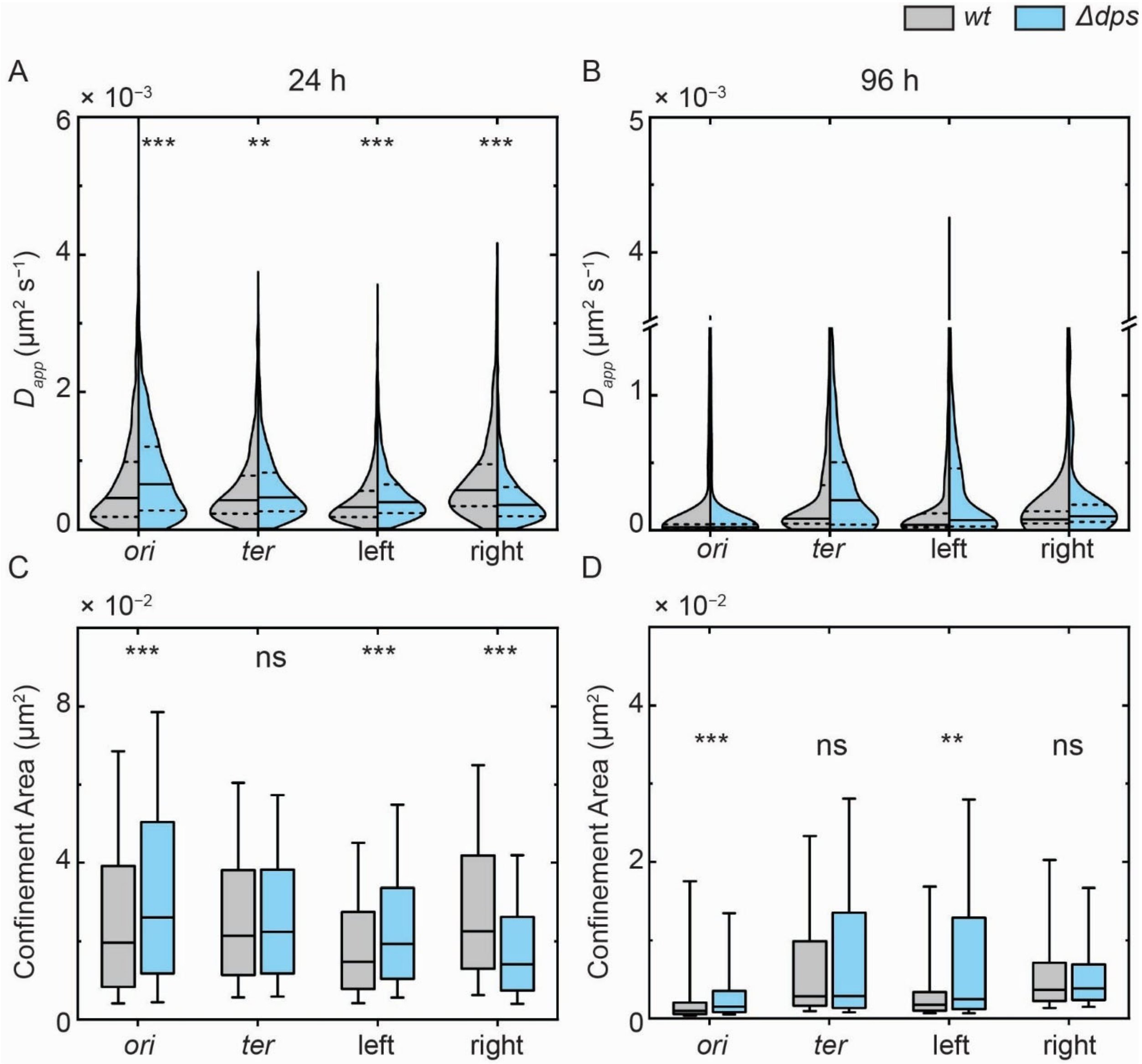

**Figure S12. A second replicate measurement of chromosomal locus dynamics at 24 and 96 h in *wt* and *Δdps* cells with only one ParB-YGFP focus with 2-s time-lapse imaging. (A)** Diffusion coefficient distribution of loci within the *ori*, *ter*, left and right macrodomains after 24 h of incubation. Dashed lines are the first and third quartiles of each distribution and solid lines are the median. $n \geq 900$ ParB-YGFP foci for each distribution. **(B)** Same as panel **A**, but for cells incubated 96 h. $n \geq 1000$ foci for the *ori, ter*, and left macrodomains and $n \geq 100$ for the right macrodomain. **(C)** Confinement area of the four loci from *wt* and *Δdps* cells at 24 h. The box length is the interquartile range, the solid line is the median and the whiskers cover the 10 – 90% range of each distribution. **(D)** Same as panel **C**, but for cells at the 96 h timepoint.

Statistical analysis was performed as described for cells with two ParB-YGFP foci. *: distributions determined to be significantly different ($p < 0.05$), **: $p < 0.01$, ***: $p < 0.001$.

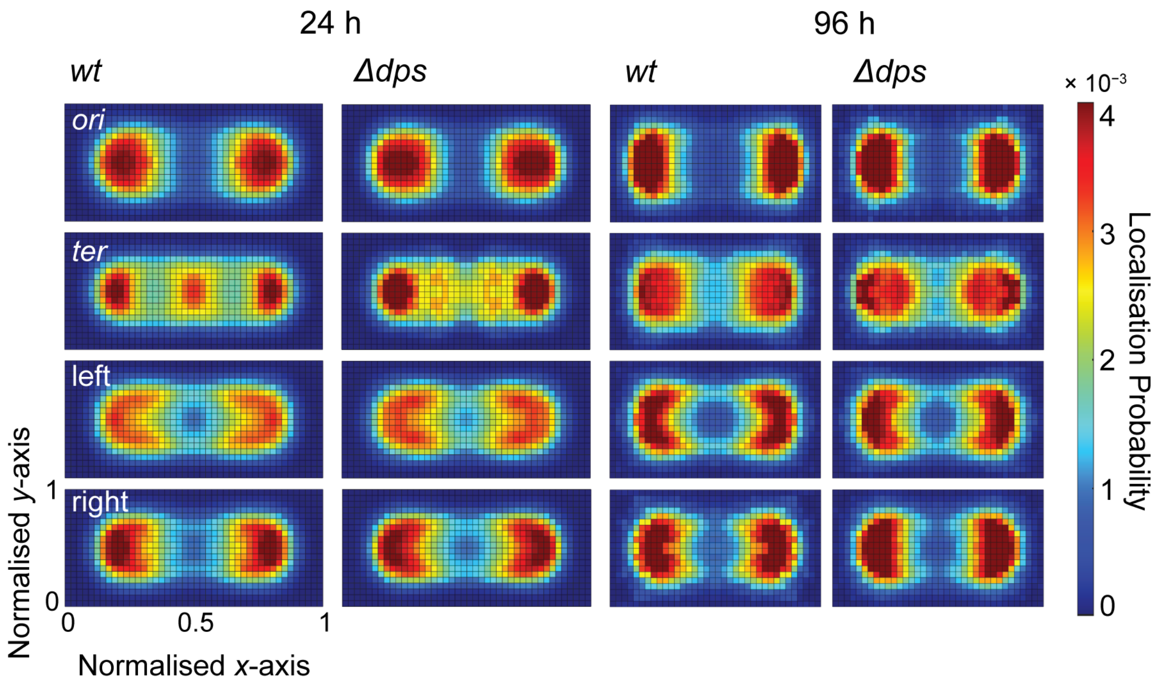

**Figure S13. Normalised heatmaps of ParB-YGFP foci localisations for four different loci located in the *ori*, *ter*, left and right macrodomains.** Dps does not significantly impact the spatial arrangement of the loci at either 24 or 96 h. Each map is constructed using data from $n \geq 1000$ cells with two ParB-YGFP foci per cell.

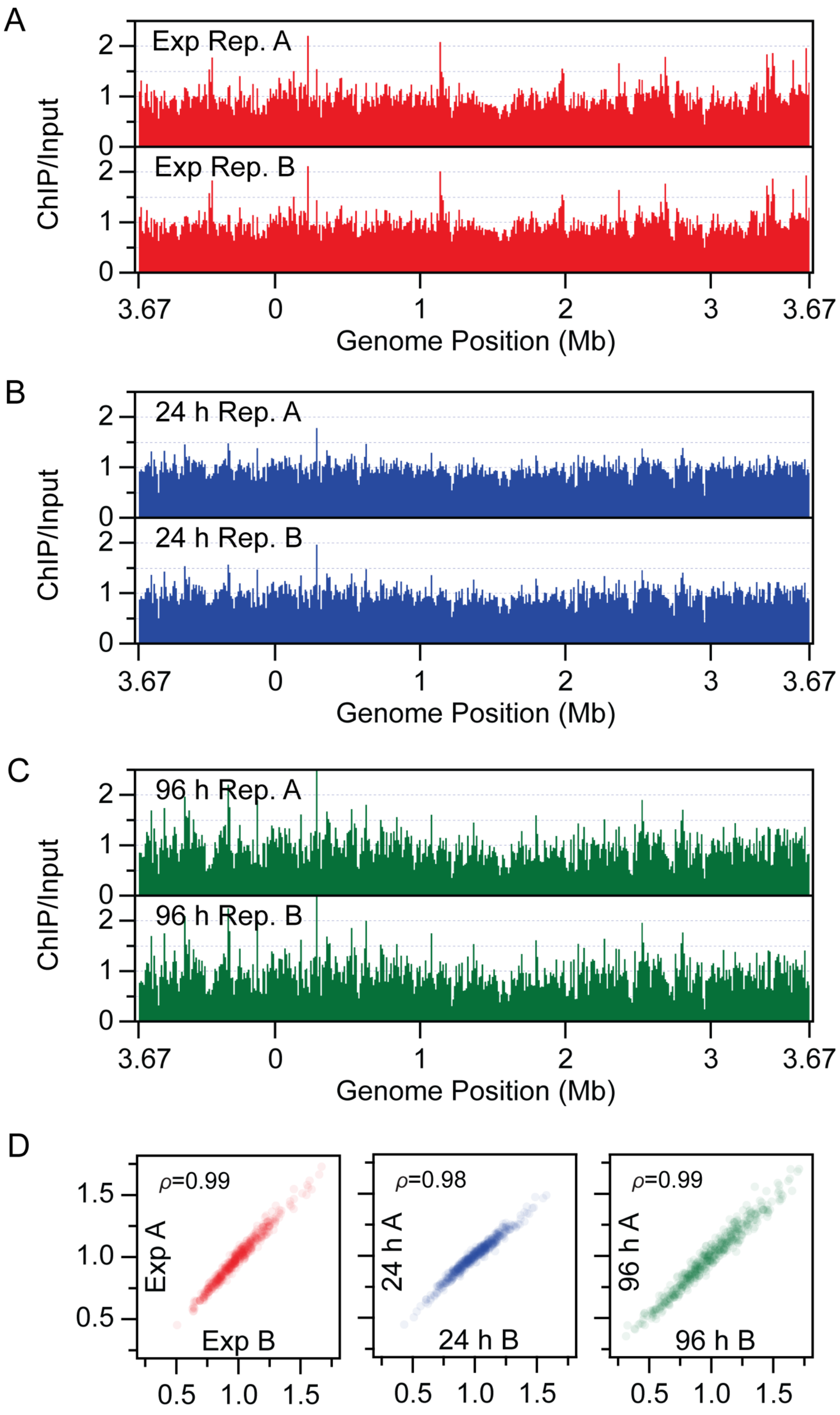

A
Exp Rep. A
Exp Rep. B
ChIP/Input
Genome Position (Mb)
3.67
0
1
2
3
3.67
B
24 h Rep. A
24 h Rep. B
ChIP/Input
Genome Position (Mb)
C
96 h Rep. A
96 h Rep. B
ChIP/Input
Genome Position (Mb)
D
$\rho$=0.99
Exp A
Exp B
$\rho$=0.98
24 h A
24 h B
$\rho$=0.99
96 h A
96 h B
0.5
1.0
1.5

**Figure S14. Biological replicates of genome-wide binding profiles of Dps. (A-C)** Biological replicates of anti-Dps ChIP-seq for each time point. ChIP-seq was performed on *wt E. coli* W3110 at **(A)** exponential phase, **(B)** stationary phase, and **(C)** deep stationary phase. The enrichment of Dps was obtained by normalizing the sequencing reads at each base pair to the total number of reads and calculating the ratio of ChIP/input. The genome was plotted using 10 kb bins and oriented to start at *oriC*. **(D)** Correlation between ChIP-seq replicates. $\rho$ is the Pearson correlation coefficient.

**Table S1. Strains used in this study**

| Strain | Genotype | Reference | Figures |
|---|---|---|---|
| cWX2782 | W3110 *hupA::PAmCherry FRT-cam-FRT* | This study | 2, S1, S2, S3, S4 |
| cWX3017 | W3110 *hupA::PAmCherry FRT-cam-FRT, Δdps kan* | This study | 2, S1, S3, S4 |
| LAM045 | W3110 *Δdps FRT-kan-FRT, pBad42 I3-01-sfgfp, spec* | This study | 3ACDE |
| LAM056 | W3110 *pBad42 I3-01-sfgfp, spec* | This study | 3ACDE |
| LAM061 | W3110 *pBad42 RBS PAmCherry-ecoRIR[E111Q], spec* | This study | 3FH, S6 |
| LAM064 | W3110 *Δdps FRT-kan-FRT, pBad42 RBS PAmCherry-ecoRIR[E111Q], spec* | This study | 3FH, S6 |
| cWX2704 | W3110 *ydeA (ter) PdnaA ygfp-ParB$^{MT1}$-parS$^{MT1}$ FRT* | This study | 4DEFGI, S7, S9, S10 |
| cWX2705 | W3110 *yfeS (left) PdnaA ygfp-ParB$^{MT1}$-parS$^{MT1}$ FRT* | This study | 4DEFGI, S7, S9, S10 |
| cWX2707 | W3110 *nlpE (right) PdnaA ygfp-ParB$^{MT1}$-parS$^{MT1}$ FRT* | This study | 4DEFGI, S7, S8, S9, S10 |
| cWX2716 | W3110 *yidA(ori) PdnaA ygfp-ParB$^{MT1}$-parS$^{MT1}$ FRT* | This study | 4ADEFGHI, S7, S9, S10 |
| cWX2724 | W3110 *ydeA (ter) PdnaA ygfp-ParB$^{MT1}$-parS$^{MT1}$ FRT, Δdps kan* | This study | 4DEFGJ, S7, S9, S10 |
| cWX2726 | W3110 *yfeS (left) PdnaA ygfp-ParB$^{MT1}$-parS$^{MT1}$ FRT, Δdps kan* | This study | 4DEFGJ, S7, S9, S10 |

| | | | |
|---|---|---|---|
| cWX2728 | W3110 *nlpE (right) PdnaA ygfp-ParB*$^{MT1}$*-parS*$^{MT1}$ *FRT, Δdps kan* | This study | 4DEFGJ, S7, S8, S9, S10 |
| cWX2730 | W3110 *yidA (ori) PdnaA ygfp-ParB*$^{MT1}$*-parS*$^{MT1}$ *FRT, Δdps kan* | This study | 4DEFGJ, S7, S9, S10 |
| W3110 | *E. coli* K-12 strain W3110 (CGSC no. 4474) *F-, λ-, IN(rrnD-rrnE)1, rph-1* | (10) | 5, 6, S11 |
| cWX2979 | W3110 (CGSC no. 4474) *Δdps FRT-kan-FRT* | This study | 6AEFG |
| cWX2695 | W3110, *rpoC::rpoC-PAmCherry FRT-kan-FRT* | This study | S5 |
| cWX2732 | W3110, *rpoC::rpoC-PAmCherry FRT no ab, Δdps FRT-kan-FRT* | This study | S5 |
| LAM059 | W3110 *pBad42 RBS ecoRIR, spec* | This study | S6 |
| LAM062 | W3110 *Δdps FRT-kan-FRT, pBad42 RBS ecoRIR, spec* | This study | S6 |
| LAM060 | W3110 *pBad42 RBS PAmCherry-ecoRIR, spec* | This study | S6 |
| AM10 | BL21(DE3) *pLysS pET17b dps, amp cm* | (21) | S6 |
| **Strains used to build strains** | | | |
| TB10 | *rph*-1 *ilvG rfb-50 λΔcro-bio nad::Tn10* | (12) | |
| AM257 | W3110 *Δdps kan* | This study | |
| hns-PAmCherry-cam | BW25993 *hupA::PAmCherry FRT-cam-FRT* | (11) | |
| cWX2565 | W3110 *yidA(ori) PdnaA ygfp-ParB*$^{MT1}$*-parS*$^{MT1}$ *FRT-kan-FRT* | This study | |
| cWX2610 | W3110, *ydeA (ter) PdnaA ygfp-ParB*$^{MT1}$*-parS*$^{MT1}$ *FRT-kan-FRT* | This study | |
| cWX2621 | W3110 *yfeS (left) PdnaA ygfp-ParB*$^{MT1}$*-parS*$^{MT1}$ *FRT-kan-FRT* | This study | |
| cWX2623 | W3110 *nlpE (right) PdnaA ygfp-ParB*$^{MT1}$*-parS*$^{MT1}$ *FRT-kan-FRT* | This study | |

| | | |
|---|---|---|
| cWX2564 | *TB10, yidA(ori)::PdnaA ygfp-ParB$^{MT1}$-parS$^{MT1}$ FRT Km* | This study |
| cWX2606 | *TB10, ydeA (ter) PdnaA ygfp-ParB$^{MT1}$-parS$^{MT1}$ FRT-kan-FRT* | This study |
| cWX2612 | *TB10, yfeS (left) PdnaA ygfp-ParB$^{MT1}$-parS$^{MT1}$ FRT-kan-FRT* | This study |
| cWX2614 | *TB10, nlpE (right) PdnaA ygfp-ParB$^{MT1}$-parS$^{MT1}$ FRT-kan-FRT* | This study |
| W3110 | *E. coli* K-12 strain W3110 ((Migula) Castellani and chalmers) *F-, λ-, IN(rrnD-rrnE)1, rph-1* | ATCC 27325 |
| CC253 | MG1655 *rpoC::rpoC-PAmCherry FRT-kan-FRT* | (14) |
| cWX2718 | W3110, *rpoC::rpoC-PAmCherry FRT no ab* | This study |

**Table S2. Plasmids used in this study**

| Plasmid | Description | Reference |
|---|---|---|
| pKD13 | *amp, FRT-kan-FRT* | (9) |
| pKD46 | *amp, red recombinase* | (9) |
| pCP20 | *bla, cat,* cI857 *repA*(*Ts*) $P_R$::*flp* | (13) |
| pKD4 | *amp, Ts, FRT-kan-FRT* (Addgene plasmid #45605) | (9) |
| pWX294 | pACYC origin with MCS, a*mp* | (7) |
| pWX950 | *pNPTS138 PT7strong yGFP-parB*$^{MT1}$*-parS*$^{MT1}$ *at Atu3054/ATU_RS14060* | (8) |
| pWX1016 | *pACYC T7T7 ygfp-ParB*$^{MT1}$*-parS*$^{MT1}$ *FRT-kan-FRT, amp* | This study |
| pWX1082 | *pACYC T7T7 PdnaA ygfp-ParB*$^{MT1}$*-parS*$^{MT1}$ *FRT-kan-FRT, amp* | This study |
| pWX1121 | *pACYC PdnaA ygfp-ParB*$^{MT1}$*-parS*$^{MT1}$ *FRT-kan-FRT, amp* | This study |
| pNC | *pET29b+ I3-01-sfgfp, kan* | (3) |
| pSCC2 | *pKC30, ecoRIR, ecoRIM, amp* (Addgene plasmid #39993) | (5) |
| pBad42 | *pBad42, spec* | (4) |
| pYH78 | *Ptrc99A-PAmCherry* | (6) |
| pLAM003 | *pBad42 I3-01-sfgfp, spec* | This study |
| pLAM012 | *pBad42 ecoRIR, spec* | This study |
| pLAM014 | *pBad42 RBS ecoRIR, spec* | This study |
| pLAM015 | *pBad42 RBS PAmCherry-ecoRIR, spec* | This study |
| pLAM016 | *pBad42 RBS PAmCherry-ecoRIR[E111Q], spec* | This study |

**Table S3. Oligonucleotides used in this study**

| Oligos | Sequence | Use |
|---|---|---|
| AM31 | TTAATCTCGTTAATTACTGGGACATAACATCAAGAGGATATGAAATT ATGGTGTAGGCTGGAGCTGCTTC | AM257 |
| AM32 | ACTAAAGTTCTGCACCATCAGCGATGGATTTATTCGATGTTAGACTC GATATTCCGGGGATCCGTCGACC | AM257 |
| AM34 | CGGTGCCCTGAATGAACTGC | Colony PCR |
| AM35 | CGGCCACAGTCGATGAATCC | Colony PCR |
| oWX418 | GTGAGTACTCAACCAAGTCATTCTGAG | pWX1121 |
| oWX2071 | CTCAGAATGACTTGGTTGAGTACTCAC | pWX1121 |
| oWX2217 | TCCTTCTGCTCCCTCGCTCAGTATGGACAGCAAGCGAACCGGAATTG | sequencing |
| oWX2379 | CTTTTTCCATCAGCTCTGTTACCG | sequencing |
| oWX2397 | GATGACGGTAACTACAAAACCC | sequencing |
| oWX2407 | CTCTAGATAGCGCATGCTGAATTC | pWX1016, sequencing |
| oWX2408 | GGTTATGCTAGTTATTGCTCAGCC | pWX1016 |
| oWX2426 | GGCCTTCTGCTTAGCTAGAGCGGC | sequencing |
| oWX2686 | GGACCATGGCTAATTCCCATACTAGTCCTTGGGGCCTCTAAACGGGT C | pWX1016 |
| oWX2687 | GAATTCAGCATGCGCTATCTAGAGGGAAACCGTTGTGGTCTCCCTAT AG | pWX1016 |
| oWX2688 | GGCTGAGCAATAACTAGCATAACCCGGGTGTAGGCTGGAGCTGCTT CG | pWX1016 |

| | | |
|---|---|---|
| oWX2689 | GACCCGTTTAGAGGCCCCAAGGACTAGTATGGGAATTAGCCATGGTCC | pWX1016 |
| oWX2765 | CCGCCCATTACACGGCATACAGCT | cWX2606, sequencing |
| oWX2766 | GAATTCAGCATGCGCTATCTAGAGCTATTGCGTCTGTT | cWX2606 |
| oWX2767 | GGACCATGGCTAATTCCCATTTGAAAGGCCCATTCGGGCC | cWX2606 |
| oWX2768 | TGGCCTCGGTGTACGTATTTGCCATCATGA | cWX2606, sequencing |
| oWX2769 | AAGCCAGTAAATTGATTGCCGCAA | cWX2612, sequencing |
| oWX2770 | GAATTCAGCATGCGCTATCTAGAGTTAATTTTCGTGAG | cWX2612 |
| oWX2771 | GGACCATGGCTAATTCCCATATACTGCATTTGTCGGCAGCAACA | cWX2612 |
| oWX2772 | TGGAGATGGCGAATCGTGGCGAAG | cWX2612, sequencing |
| oWX2773 | TCGAAAAAGACGGAACATGGGTGA | cWX2614, sequencing |
| oWX2774 | GAATTCAGCATGCGCTATCTAGAGTTACTGCCCCAAACTA | cWX2614 |
| oWX2775 | GGACCATGGCTAATTCCCATCCCGTCTTGAGACAGAAACA | cWX2614 |
| oWX2776 | ATAACGTTGCAGAAGCGACAGGCG | cWX2614, sequencing |
| oWX2815 | TCGCCGCTGGCCAGTGACACTCGAAGAACAGACGCAATAGCTCTAGATAGCGCATGCTGA | cWX2606 |
| oWX2816 | TAAAACGTACCATTAAAAAAGGCCCGAATGGGCCTTTCAAATGGGAATTAGCCATGGTCC | cWX2606 |
| oWX2817 | AATGATTGATGACCTTACGGCGTTTCCTCACGAAAATTAACTCTAGATAGCGCATGCTGA | cWX2612 |
| oWX2818 | GCACTTTTTTAACAGTTGTTGCTGCCGACAAATGCAGTATATGGGAATTAGCCATGGTCC | cWX2612 |

| | | |
|---|---|---|
| oWX2819 | ATTTTACCCCAACCAGGATTGCAGTAGTTTGGGGCAGTAACTCTAGATAGCGCATGCTGA | cWX2614 |
| oWX2820 | CTTCTGGCCTGTTTTGCGTTTGTTTCTGTCTCAAGACGGGATGGGAATTAGCCATGGTCC | cWX2614 |
| oWX2844 | AAATGATGACCTCGTTTCCACC | pWX1082 |
| oWX2946 | AGCTTAAAGGAGGTGGAAACATG | pWX1082 |
| oWX2947 | GTTTCCACCTCCTTTAAGCTTCCACTCGAACAAAAGTCGATAATG | pWX1082 |
| oWX2948 | GGTGGAAACGAGGTCATCATTTCAATAATTGTACACTCCGGAGTC | pWX1082 |
| oWX2981 | ACCTTGAAGATGGCGTGGCGTTTGCTATTGAGAAGTATGTGCTGAATTAACTCTAGATAGCGCATGCTGA | cWX2564 |
| oWX2982 | AATAATCAGTAAGCGGGCAAACGCGTTTATGCTGTTTGCCCGCCCACAGAATGGGAATTAGCCATGGTCC | cWX2564 |
| oWX3021 | ATGACCGTTCACATCACCATCCAG | sequencing |
| oWX3052 | CTCTAGATAGCGCATGCTGAATAATAATTGTACACTCCGGAGTC | pWX1121 |
| oWX3053 | ATTCAGCATGCGCTATCTAGAG | pWX1121 |
| LM022 | GTTTTTTTGGGCTAGCGTTTGTTTAACTTTAAGAAGGAGATATACATATGCACCACC | pLAM003 |
| LM024 | TTGCATGCCTGCAGGCAACTCAGCTTCCTTTCG | pLAM003 |
| LM045 | GTTTTTTTGGGCTAGCGATGTCTAATAAAAAACAGTCAAATAGGCTAACTGAAC | pLAM012 |
| LM046 | GTTTTTTTGGGCTAGCGATGTCTAATAAAAAACAGTCAAATAGGCTAACTGAAC | pLAM012 |
| LM050 | AGGAGGTGAAATTGCTAGCGATGTCTAATAAAAAACAG | pLAM014, pLAM015 |
| LM051 | TTGCATGCCTGCAGG | pLAM014, pLAM015 |

| | | |
|---|---|---|
| LM052 | GTTTTTTTGGGCTAGCGAGGAGGTGAAATTGCTAGC | pLAM014, pLAM015 |
| LM053 | AGGAGGTGAAATTGCTAGCGATGAGCAAGGGCGAGGAG | pLAM015 |
| LM054 | GCTACCGCTGCCGGAGCCGGAACCGGAGCCCTTGTACAGCTCGTC | pLAM015 |
| LM055 | ATTAGACATGCTGCTACCGCTGC | pLAM015 |
| LM056 | GGTAGCAGCATGTCTAATAAAAAACAGTCAAATAGGCTAAC | pLAM015 |
| LM069 | ACTTGTTGCTcagGCCAAACACC | pLAM016 |
| LM070 | ACAACTCTCCATTCACC | pLAM016 |
| LM006 | GCGTTCTGATTTAATCTGTATCAGG | Colony PCR |
| LM027 | CACACTTTGCTATGCCATAGC | Colony PCR |

**Table S4. NGS samples in this study** (reviewer token: ylynsaysfduvzmx)

| Sample name | Figure | Reference | Identifier |
|---|---|---|---|
| ChIP_anti-Dps_cWX2200_exp_rep1 | 5 | This study | GSM8885036 |
| ChIP_anti-Dps_cWX2200_24H_rep1 | 5 | This study | GSM8885034 |
| ChIP_anti-Dps_cWX2200_96H_rep1 | 5 | This study | GSM8885035 |
| ChIP_anti-mCherry_cWX2200_96H_rep1 | 5 | This study | GSM8885037 |
| WGS_cWX2200_exp_rep1_ChIPinput_for_Dps | 6 | This study | GSM8885041 |
| WGS_cWX2200_24H_rep1_ChIPinput_for_Dps | 6 | This study | GSM8885038 |
| WGS_cWX2200_96H_rep1_ChIPinput_for_Dps | 6 | This study | GSM8885039 |
| WGS_cWX2200_96H_rep2_ChIPinput_for_mCherry | 6 | This study | GSM8885040 |
| cWX2979_96H_Azure_10percent_Sau3AI_rep1 | 6AEG | This study | GSM8885046 |
| cWX2979_24H_Azure_10percent_Sau3AI_rep1 | 6AEF | This study | GSM8885045 |
| cWX2979_exp_Azure_10percent_Sau3AI_rep1 | 6A | This study | GSM8885047 |
| cWX2200_96H_Azure_10percent_Sau3AI_rep2 | 6ABDEG | This study | GSM8885043 |
| cWX2200_24H_Azure_10percent_Sau3AI_rep1 | 6ABCDEF | This study | GSM8885042 |
| cWX2200_exp_Azure_10percent_Sau3AI_rep2 | 6ABC | This study | GSM8885044 |

**Table S5. Mean nucleoid occupancies of stationary and late-stationary-phase cells quantified *via* bulk fluorescence microscopy of SYTOX green and super-resolution imaging of HUα-PAmCherry**

| Strain | Condition | Mean ± standard deviation: Nucleoid occupancy measured by bulk fluorescence | Mean ± standard deviation: Nucleoid occupancy measured by super-resolution |
|---|---|---|---|
| *wt* | 24 h | 0.57 ± 0.16 | 0.52 ± 0.05 |
| *Δdps* | 24 h | 0.56 ± 0.14 | 0.51 ± 0.04 |
| *wt* | 96 h | 0.55 ± 0.14 | 0.49 ± 0.10 |
| *Δdps* | 96 h | 0.55 ± 0.13 | 0.55 ± 0.07 |

**Supplementary Information References**
